# Biochemical and physiological characterization of *Aedes aegypti* midgut chymotrypsin

**DOI:** 10.1101/2024.12.31.630969

**Authors:** Abigail G. Ramirez, Jun Isoe, Mateus Sá M Serafim, Daniel Fong, My Anh Le, James T. Nguyen, Olive E. Burata, Rachael M. Lucero, Alberto A. Rascón

## Abstract

The *Aedes aegypti* mosquito is a vector of dengue, Zika, and chikungunya. The mosquito’s reliance on blood facilitates the transmission of these viral pathogens to humans. Digestion of blood proteins depends on the biphasic expression of serine proteases, with trypsin-like activity contributing to most of the activity in the midgut. Other proteases found (serine collagenase- and chymotrypsin-like) are thought to contribute to digestion, but their roles are largely understudied. Thus, elucidating the activity and specific roles of all midgut proteases will help understand the complexity of the digestion process and help validate them as potential targets for the development of a new vector control strategy. Herein, we focused on characterizing the activity profile and role of *Ae. aegypti* chymotrypsin (AaCHYMO). Knockdown studies resulted in elimination and significant reduction of chymotrypsin-like activity in blood fed midgut extracts, while *in vitro* fluorescent and blood protein digestion assays revealed important substrate specificity differences. Interestingly, knockdown of AaCHYMO did not impact fecundity, indicating the presence of an intricate network of proteases working collectively to degrade blood proteins. Further, knockdown of the ecdysone receptor (EcR) led to a decrease in overall AaCHYMO expression and activity in the mosquito, which may play an important regulatory role.

## INTRODUCTION

The *Aedes aegypti* mosquito relies on the digestion of blood meal proteins by midgut proteases to successfully complete the gonotrophic cycle and lay eggs. Two major classes of proteolytic enzymes responsible for this activity are serine-like endopeptidases (mainly trypsin-, chymotrypsin-, and serine collagenase-like) and exopeptidases that are released into the midgut lumen after imbibing a blood meal ^1–3^. Blood meal digestion is biphasic with trypsin-like activity responsible for most, if not all, proteolytic activity found in both the early phase (beginning immediately after a blood meal) and the late phase (beginning around 12 to 18 h post-blood meal (PBM)) ^1–3^. Because of this, *Ae. aegypti* midgut blood meal digestion studies have mainly focused on trypsin-like proteases ^1,3–5^, with the contribution of chymotrypsin-like proteases being minimal or relatively unknown. Since 1997, only three chymotrypsin-like proteases have been described in the literature. The first of these enzymes, AaCHYMO, led to an increase in protein expression and activity in mosquito midgut extracts, only after a blood meal, suggesting a role in digestion ^6^. The second chymotrypsin-like protease (Juvenile Hormone Associated 15, JHA15) was identified in 2008, and initial studies showed that the knockdown of the protease leads to no phenotypic changes in the mosquito despite being present in the midgut during the early phase portion of blood meal digestion ^7^. The third chymotrypsin-like protease (chymotrypsin II, AaCHYMO II) was identified in 2023, and was shown to be present in both larvae and adult female mosquitoes, but its role in blood meal digestion was not explored ^8^.

Due to the *Ae. aegypti* mosquito reliance on blood to fuel the gonotrophic cycle, elucidating the activity of all midgut proteases will help better understand the complexity of the blood meal digestion process. In doing so, we can validate these proteolytic enzymes as potential inhibitor targets ^1,3–5^. The *Aedes* mosquito species’ (*Ae. aegypti* and *Ae. albopictus*) reliance on blood has facilitated the spread of arthropod-borne viruses (*i.e.*, arboviruses), such as dengue (DENV), Zika (ZIKV), and chikungunya (CHIKV) to uninfected human hosts ^9^. The prevalence of ZIKV has been on the rise since 2007, with a major outbreak in the United States in 2015, which was denoted by severe clinical manifestations (organ failure), birth defects (microcephaly), fetal death, and Guillain-Barré syndrome ^10^. Although ZIKV, DENV, and CHIKV are more prevalent in tropical settings, due to the ever-changing climate, *Ae. aegypti* and *Ae. albopictus* have been present in Florida and Texas ^11,12^. In addition, low-level locally acquired DENV and CHIKV infections have been reported and ongoing in these regions, and may become more susceptible to viral transmission, leading to a permanent establishment of locally acquired infections ^11,12^.

Fortunately, insecticides remain the only successful vector control strategy, but even then, the heavy use of commercially available pesticides has led to resistance in *Ae. aegypti* mosquitoes ^13,14^. Biological methods, such as the use of sterile- or *Wolbachia*-infected mosquitoes, have shown promise in suppressing the mosquito population and viral replication, but logistical issues in production and release of these mosquitoes have limited their use ^13,15,16^. Given these limitations, targeting blood meal protein digestion may be an alternative approach to mosquito population control. After imbibing a blood meal, midgut proteases are expressed and released into the midgut lumen which then process blood proteins into the necessary nutrients used by the mosquito. In certain cases, when midgut protease expression is knocked down (as is the case with *Ae. aegypti* Serine Protease VI (*AaSPVI*), *AaSPVII*, and Late Trypsin (*AaLT*)) fecundity is affected leading to ∼22.9% overall reduction in egg production ^1^. However, the targeting of midgut proteases may be challenging given the complex network of other endo- and exopeptidases that are released ^1–3,17^.

Recently, we determined the global proteolytic profile of *Ae. aegypti* midgut tissue extracts from sugar and blood fed mosquitoes using Multiplex Substrate Profiling by Mass Spectrometry (MSP-MS), revealing the synergistic importance of proteolytic activity on the digestion of blood meal proteins ^3^. Surprisingly, AaCHYMO alone cleaved 22% (32/146) of the peptide bonds in the peptide library suggesting a potentially important role in blood meal protein digestion. Thus, in the present study we focused on determining the activity profile and role of AaCHYMO and ascertain its significance in the *Ae. aegypti* mosquito gonotrophic cycle. Knockdown of AaCHYMO in the mosquito led to elimination and significant reduction in chymotrypsin-like activity in midgut extracts. In addition, the knockdown of the ecdysone receptor (EcR), a key nuclear receptor that plays an important role in the mosquito gonotrophic cycle ^18^, led to an effect on AaCHYMO expression and chymotrypsin-like activity in mosquito midgut extracts. Further, recombinant AaCHYMO was expressed, purified, and activated for *in vitro* fluorescent and blood protein digestion activity assays, and compared to human and bovine chymotrypsin. Results reveal important substrate specificity differences, especially in the degradation of human blood meal proteins. Taken together, understanding the role of AaCHYMO and other understudied midgut proteases will aid in validating these enzymes as potential targets for the development of inhibitors and a new vector control strategy.

## RESULTS

### Proteolytic activity assays indicate specific substrate preferences between mosquito AaCHYMO, human and bovine chymotrypsin

Active recombinant AaCHYMO activity was tested against several commercially available chymotrypsin substrates (**Supplementary Table 1**) and fit to the Michaelis-Menten (MM) equation (representative MM plots are shown in **Supplementary Figure 1**). Out of the 14 chymotrypsin substrates tested, MeO-Suc-Arg-Pro-Tyr-AMC resulted in the best kinetic parameters with *k_cat_* = 14.2 sec^-1^, *K_m_* = 9.52 μM, and *k_cat_/K_m_* = 1,481,520 M^-1^ s^-1^ (**Table 1**). For comparison, commercially available human and bovine chymotrypsin were also tested and found to have higher overall kinetic parameters than AaCHYMO for this substrate and the other commercially available chymotrypsin substrates. In some cases, *k_cat_/K_m_* values for human and bovine chymotrypsin were more than 26- and 6000-fold higher than AaCHYMO, respectively (**Table 1**). The worst commercially available substrate processed by AaCHYMO (Suc-Leu-Leu-Val-Tyr-AMC, *k_cat_* = 0.063 sec^-1^, *K_m_* = 6.72 μM, *k_cat_/K_m_* = 9,410 M^-1^ s^-1^) was the overall most processed by both human (*k_cat_* = 597 sec^-1^, *K_m_* = 6.65 μM, *k_cat_/K_m_* = 89,814,954 M^-1^ s^-1^) and bovine chymotrypsin (*k_cat_* = 580.3 sec^-1^, *K^m^* = 10.67 μM, *k^cat^/K^m^* = 54,389,253 M^-1^ s^-1^). Interestingly, human chymotrypsin did not readily process chymotrypsin substrates with a Leu at the P1 site, while AaCHYMO and bovine chymotrypsin did, with AaCHYMO having an overall higher specificity constant (*k_cat_/K_m_*) when Leu is at the P1 site (**Table 1**). In addition, two trypsin-like substrates (Boc-Val-Leu-Lys-AMC and Z-Arg-Arg-AMC) were tested, and only human chymotrypsin readily processed these substrates.

**Table 1.** Kinetic Parameters Determined for AaCHYMO Using Commercially Available Chymotrypsin- and AaCHYMO-Specifically Designed AMC Fluorescent Substrates Compared to Human and Bovine Chymotrypsin.

| Commercially Available | AaCHYMO |  |  |  | Human Chymotrypsin |  |  |  | Bovine Chymotrypsin |  |  |  | Human/AaCHYMO | Bovine/AaCHYMO |
| --- | --- | --- | --- | --- | --- | --- | --- | --- | --- | --- | --- | --- | --- | --- |
| Substrate | $k_{cat}$ (s<sup>-1</sup>) | $K_m$ (μM)* | $k_{cat}/K_m$ (M<sup>-1</sup> s<sup>-1</sup>) | Specific Activity<br>μmol min<sup>-1</sup> (mg protein)<sup>-1</sup> | $k_{cat}$ (s<sup>-1</sup>) | $K_m$ (μM)* | $k_{cat}/K_m$ (M<sup>-1</sup> s<sup>-1</sup>) | Specific Activity<br>μmol min<sup>-1</sup> (mg protein)<sup>-1</sup> | $k_{cat}$ (s<sup>-1</sup>) | $K_m$ (μM)* | $k_{cat}/K_m$ (M<sup>-1</sup> s<sup>-1</sup>) | Specific Activity<br>μmol min<sup>-1</sup> (mg protein)<sup>-1</sup> | ratio | ratio |
| Suc-Ala-Ala-Pro-Phe-AMC | 4.81 | 9.72 | 4949578 | 11.0 | 608.7 | 114.2 | 5329831 | 1460.8 | 832 | 18.77 | 44326052 | 1997 | 1.08 | 9.0 |
| Suc-Ala-Ala-Phe-AMC | 0.92 | 83.16 | 11087 | 2.10 | 8.38 | 30.0 | 279724 | 20.1 | 100.3 | 48.44 | 2069915 | 240.6 | 25.2 | 186.7 |
| MeO-Suc-Arg-Pro-Tyr-AMC | 14.1 | 9.52 | 1481520 | 32.1 | 1597 | 12.3 | 129915921 | 3832 | 3146 | 8.014 | 392563015 | 7550 | 87.7 | 265.0 |
| Suc-Leu-Leu-Val-Tyr-AMC | 0.06 | 6.72 | 9410 | 0.14 | 597 | 6.6 | 89814954 | 1432.8 | 580.3 | 10.67 | 54389253 | 1393 | 9545 | 5780 |
| Suc-Arg-Pro-Phe-His-Leu-Leu-Val-Tyr-MCA | 0.61 | 1.54 | 394925 | 1.38 | 37.4 | 1.2 | 31562676 | 89.8 | 6.82 | 0.57 | 11959064 | 16.4 | 79.9 | 30.3 |
| Suc-Leu-Tyr-AMC | 0.07 | 7.57 | 8712 | 0.15 | 5.16 | 10.6 | 485732 | 12.4 | 8.58 | 16.29 | 526499 | 20.6 | 55.8 | 60.4 |
| Suc-Ile-Ile-Trp-MCA | 5.14 | 61.68 | 83279 | 11.7 | 794 | 12.2 | 65188834 | 1905.6 | 7893 | 13.37 | 590376465 | 18944 | 782.8 | 7089 |
| Ac-Ala-Asn-Trp-AMC | 2.69 | 33.31 | 80636 | 6.12 | 425 | 25.1 | 16932271 | 1020 | 255.9 | 56.16 | 4556624 | 614.2 | 210.0 | 56.5 |
| Z-Leu-Leu-Leu-MCA | 0.03 | 0.13 | 204048 | 0.062 | - | - | - | - | - | - | - | - | - | - |
| Ac-Pro-Ala-Leu-AMC | 0.02 | 9.00 | 1818 | 0.037 | - | - | - | - | 0.0061 | 7.63 | 801 | 0.014664 | - | 0.44 |
| Z-Gly-Gly-Leu-AMC | - | - | - | - | - | - | - | - | - | - | - | - | - | - |
| Z-Val-Lys-Met-MCA | 0.07 | 1.21 | 60149 | 0.166 | 0.41 | 6.4 | 64896 | 0.991 | 6.32 | 6.06 | 1043215 | 15.2 | 1.08 | 17.3 |
| Boc-Val-Leu-Lys-AMC | - | - | - | - | 5.17 | 225.5 | 22942 | 12.4 | - | - | - | - | - | - |
| Z-Arg-Arg-AMC | - | - | - | - | 4.79 | 568.2 | 8424 | 11.5 | - | - | - | - | - | - |
| AaCHYMO-Specifically Designed | AaCHYMO |  |  |  | Human Chymotrypsin |  |  |  | Bovine Chymotrypsin |  |  |  | Human/AaCHYMO | Bovine/AaCHYMO |
| Substrate | $k_{cat}$ (s<sup>-1</sup>) | $K_m$ (μM)* | $k_{cat}/K_m$ (M<sup>-1</sup> s<sup>-1</sup>) | Specific Activity<br>μmol min<sup>-1</sup> (mg protein)<sup>-1</sup> | $k_{cat}$ (s<sup>-1</sup>) | $K_m$ (μM)* | $k_{cat}/K_m$ (M<sup>-1</sup> s<sup>-1</sup>) | Specific Activity<br>μmol min<sup>-1</sup> (mg protein)<sup>-1</sup> | $k_{cat}$ (s<sup>-1</sup>) | $K_m$ (μM)* | $k_{cat}/K_m$ (M<sup>-1</sup> s<sup>-1</sup>) | Specific Activity<br>μmol min<sup>-1</sup> (mg protein)<sup>-1</sup> | ratio | ratio |
| Ac-Ala-His-Val-Phe-AMC | 26.9 | 8.11 | 33173056 | 61.3 | 83.8 | 30.0 | 2797242 | 201.2 | 339.7 | 31.0 | 10974690 | 815.2 | 0.084 | 0.331 |
| Ac-Ala-His-Leu-Phe-AMC | 10.6 | 4.40 | 2419697 | 24.3 | 23.4 | 13.7 | 1706754 | 56.2 | 40.1 | 14.1 | 2849692 | 96.2 | 0.705 | 1.18 |
| Ac-Ala-His-Val-Tyr-AMC | 0.817 | 7.04 | 116115 | 1.86 | 8.4 | 25.9 | 325097 | 20.2 | 8.58 | 23.9 | 359737 | 20.6 | 2.80 | 3.10 |
| Ac-Ala-His-Leu-Tyr-AMC | 6.07 | 1.24 | 4888531 | 13.8 | 248.3 | 12.7 | 19548556 | 595.8 | 642.7 | 19.8 | 32408808 | 1542.4 | 4.00 | 6.63 |
| Ac-Ala-His-Thr-Trp-AMC | 0.072 | 7.30 | 9815 | 0.163 | 190.0 | 25.1 | 7583500 | 455.9 | 339.7 | 19.9 | 17094447 | 815.2 | 772.6 | 1741.7 |
| Ac-Ala-His-Asp-Trp-AMC<sup>†</sup> | - | - | - | - | 48.5 | 14.8 | 3285908 | 116.4 | 272.7 | 20.2 | 13525132 | 654.4 | - | - |
| Ac-Ala-His-Val-Trp-AMC | 5.51 | 6.32 | 872249 | 12.6 | 438.7 | 18.9 | 23222163 | 1052.8 | 1839.7 | 15.3 | 120318291 | 4415.2 | 26.6 | 137.9 |
| Ac-Ala-His-Val-Leu-AMC | 5.42 | 3.95 | 1372350 | 12.3 | 1.26 | 33.1 | 38176 | 3.03 | 2.30 | 31.9 | 72076 | 5.51 | 0.028 | 0.053 |
| Ac-Pro-His-Val-Met-AMC | 133.1 | 10.7 | 12482802 | 303.4 | 2.9 | 18.3 | 157534 | 6.90 | 4.19 | 11.3 | 371420 | 10.1 | 0.013 | 0.030 |
| Ac-Ala-His-Val-Met-AMC | 2.83 | 4.30 | 658452 | 6.46 | 8.61 | 43.8 | 196589 | 20.7 | 8.93 | 29.7 | 300460 | 21.4 | 0.299 | 0.456 |
| *$K_m$ and $v_{max}$ determined with the Michaelis-Menten non-linear regression analysis tool at a 95% Confidence Interval (CI) using GraphPad Prism. | | | | | | | | | | | | | | |
| - Negligible activity detected under any of the conditions tested, exceeded lower level of detection for AMC >1 nM. |  |  |  |  |  |  |  |  |  |  |  |  |  |  |
| † Non-preferred amino acid at the P2 site |  |  |  |  |  |  |  |  |  |  |  |  |  |  |

Given the differences in chymotrypsin-like activity observed, we ordered AaCHYMO-specific fluorescent AMC substrates with preferred amino acids at the P1 and P2-P4 sites, based on previously characterized proteolytic profiles of AaCHYMO ^3^. Using these specifically designed substrates, we determined AaCHYMO kinetic parameters and compared them to human and bovine chymotrypsin (**Table 1**). Amino acid residues at the P1 position influenced the activity of AaCHYMO resulting in the protease better processing of AaCHYMO-specific AMC substrates. For example, when Phe and Met are at the P1 site, AaCHYMO had an overall better specificity constant (*k_cat_/K_m_*) than human and bovine chymotrypsin. Conversely, AaCHYMO was not as efficient as the mammalian chymotrypsin proteases when Trp or Tyr are at P1 (**Table 1**). The only exception where AaCHYMO had no detectable activity from one of the AaCHYMO designed substrates was with Ac-Ala-His-Asp-Trp-AMC. In this case, the substrate was designed with a non-preferred amino acid at the P2 position (aspartate) ^3^ to determine the importance of amino acids beyond the P1 site. Human and bovine chymotrypsin readily cleaved the substrate, having *k_cat_*/*K_m_* values greater than 3×10^6^ M^-1^ s^-1^.

### Western blot analysis of AaCHYMO expression, and proteolytic assays reveal activity in previously undetected mosquito extracts and tissues

We aimed to detect if AaCHYMO would be present in blood fed midgut extracts (with and without the food bolus) and at different life stages. First, AaCHYMO was detected in midgut extracts prepared without the food bolus (midgut epithelial cells only) (**Figure 1A**). In sugar-fed (SF) samples, AaCHYMO is clearly present, followed by detection in the 3 to 24 h PBF samples, absent at 36 and 48 h, and relative protease expression observed once again at 72 h PBF. Using the midgut epithelial cell samples only, relative AaCHYMO protein levels were quantified using ImageJ (**Figure 1B**). For comparison, midgut extracts with both the midgut epithelial cells and food bolus were analyzed (**Figure 1C**). In these samples, AaCHYMO is detected at all the time points tested, with minimal detection in SF and at 72 h PBF. Due to the presence of AaCHYMO in the epithelial cells and food bolus, these samples were used to confirm the presence of chymotrypsin-like activity in mosquito midgut tissue extracts. Activity was confirmed using the optimal commercially available chymotrypsin substrate (MeO-Suc-Arg-Pro-Tyr-AMC, see **Supplementary Figure 1B** and **Table 1**). Interestingly, no detectable chymotrypsin-like activity is observed in sugar-fed and 72 h PBF midgut extract samples, indicating that the species present is the zymogen form of AaCHYMO (**Figure 1D**). An increase in chymotrypsin-like activity observed at 12, 24, and 36 h PBF correlates with active AaCHYMO, with minimal activity observed at 48 h PBF. Second, to confirm that AaCHYMO-like activity is only found in the midgut of the female mosquito, different tissues were also processed and tested. The pupae, male adult mosquitoes, female carcasses (without the midgut), and the midgut without the food bolus were found to have minimal detection of chymotrypsin-like activity when compared to the larvae and the food bolus (**Figure 1E**), as well as the midgut extracts at the different timepoints PBF (**Figure 1C**). This indicates that AaCHYMO is localized to the midgut and activated only after a blood meal is ingested.

**Figure 1.**
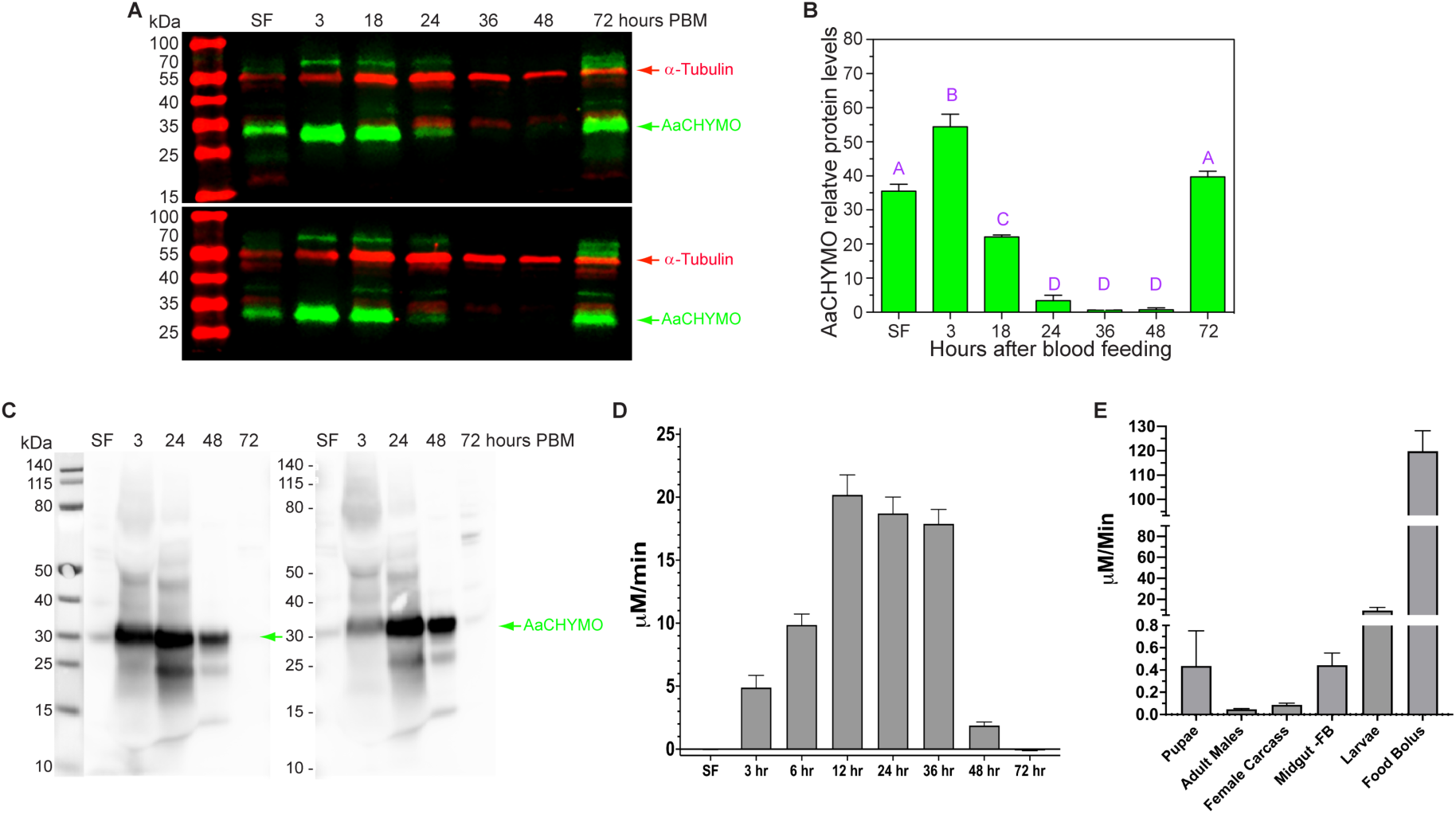
Western blot (WB) analysis and chymotrypsin-like activity detection in different *Ae. aegypti* mosquito tissues. **A.** Representative fluorescent WB (two independent sets) of sugar (SF) and blood fed midgut epithelial cell samples without the food bolus detecting the presence of AaCHYMO at the time points indicated post blood feeding (PBF). α-tubulin was used as an internal control. **B.** Relative AaCHYMO protein levels detected in SF and blood fed mosquito midgut epithelial cell samples. Experiments were conducted from 3 biological cohorts using 5 to 10 mosquitoes. **C.** Representative chemiluminescent WB (two independent sets) of sugar- and blood-fed midgut extracts that contain the midgut epithelial cells plus the food bolus (0.5 midgut equivalents were used for analysis). These are representative samples that were used for activity assays. **D.** Chymotrypsin-like activity detected in midgut extracts with the food bolus using the MeO-Suc-Arg-Pro-Tyr-AMC substrate, which corresponds to the presence of active AaCHYMO in the 3 h to 48 h PBF samples. **E.** Detection of minimal chymotrypsin-like activity in pupae, adult male mosquitoes, female carcasses (without the midgut), and midgut tissues without the food bolus (-FB). Higher chymotrypsin-like activity was detected in larvae and the food bolus when compared to the other tissues. Four sets of three individual mosquitoes used for the assays in D and E.

### Knockdown studies of AaCHYMO led to the detection of residual chymotrypsin-like activity and no effect on fecundity

We proceeded to determine the overall activity contribution of AaCHYMO in the mosquito midgut. After knockdown of the gene, AaCHYMO mRNA and protein expression were eliminated at 3 h and 18 h PBF (**Figures 2A-2D**). When comparing FLUC *vs* AaCHYMO knocked down midgut tissue extracts, complete elimination of chymotrypsin-like activity is observed from 3 h to 24 h PBF, and a significant reduction in overall activity is observed at 36 h PBF (****p < 0.0001) and 48 h PBF (**p < 0.05) (**Figures 2E-2F**). Since chymotrypsin-like activity was not eliminated at 36 h and 48 h PBF, we decided to perform a double knockdown of AaCHYMO and the late phase protease, AaLT (**Figure 3**). Recent MSP-MS profiling of AaLT revealed that the protease is not a trypsin but rather has more chymotrypsin- and serine collagenase-like specificity ^3^. This would help identify if the remaining activity is coming from AaLT or a different protease. Although both proteases were knocked down, verified via WB (**Figures 3A** and **3C**), and both protein (**Figures 3B** and **3D**) and mRNA (**Figures 3E-3G**) expression levels quantified, there were no significant changes in chymotrypsin-like activity using the MeO-Suc-Arg-Pro-Tyr-AMC substrate at the 36 h and 48 h PBF timepoints (**Figure 3I**) indicating that another chymotrypsin-like protease may be responsible for the residual activity observed.

**Figure 2.**
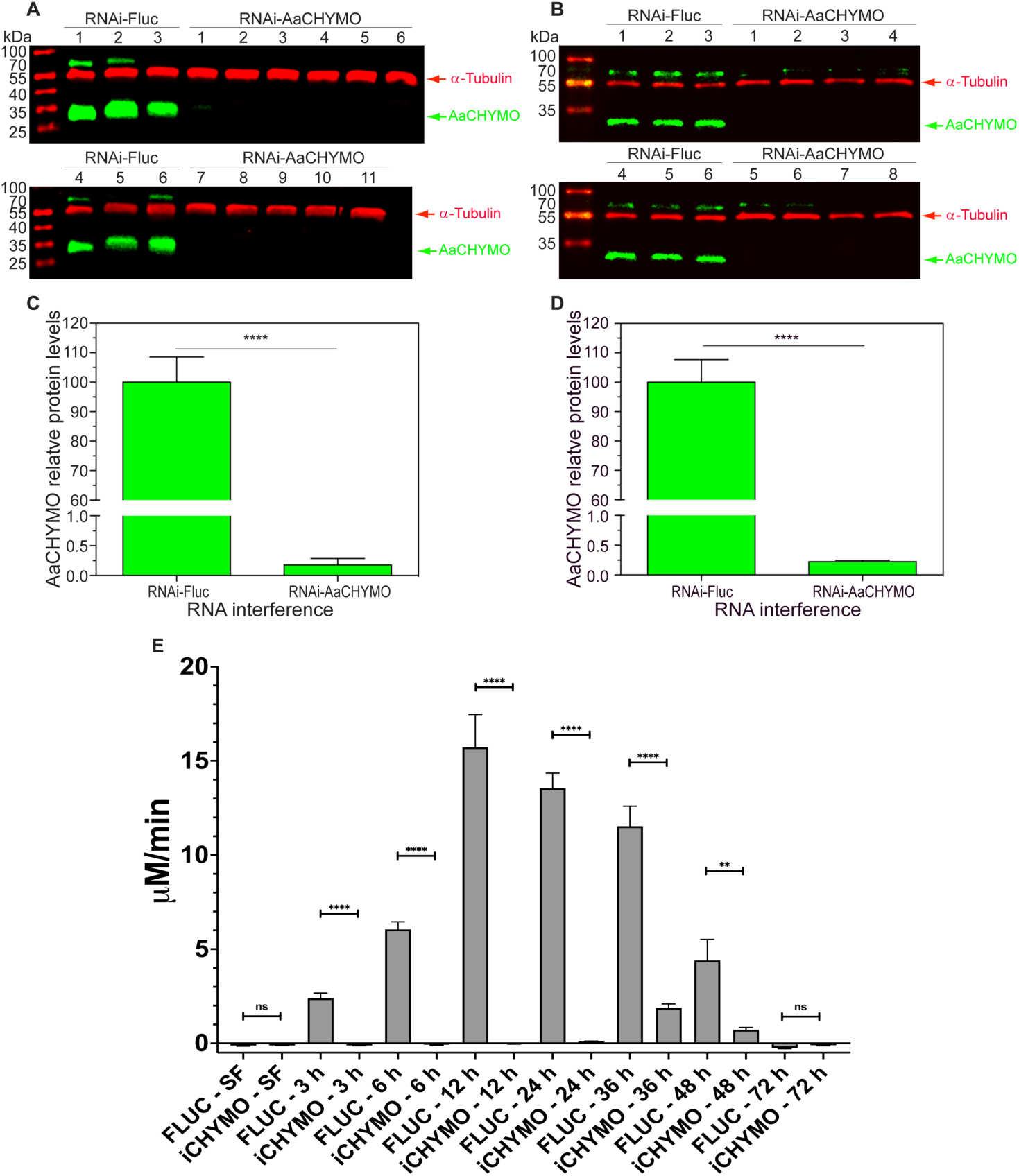
RNAi knockdown efficiency and chymotrypsin-like activity determination in individual *Ae. aegypti* mosquito midgut tissue extracts. Representative western blot of mosquito midgut extracts at 3 h PBM (**A**) and 18 h PBM (**B**). Quantification of AaCHYMO protein expression (**C**) and quantification of mRNA levels via qPCR (**D**). Data was obtained from individual midgut of 12 mosquitoes. Data are presented as mean ± SEM. Statistical significance is represented by stars above each comparison (unpaired Student’s t-test; **** p < 0.0001 compared to the RNAi-FLUC control samples). Detection of chymotrypsin-like activity in FLUC *vs* RNAi AaCHYMO mosquito midgut extracts using the MeO-Suc-Arg-Pro-Tyr-AMC substrate (**E**). Activity is eliminated in the 3 h to 24 h PBF samples, with an overall significant reduction in chymotrypsin-like activity at 36 h PBM (****p < 0.0001) and at 48 h PBM (**p < 0.05).

**Figure 3.**
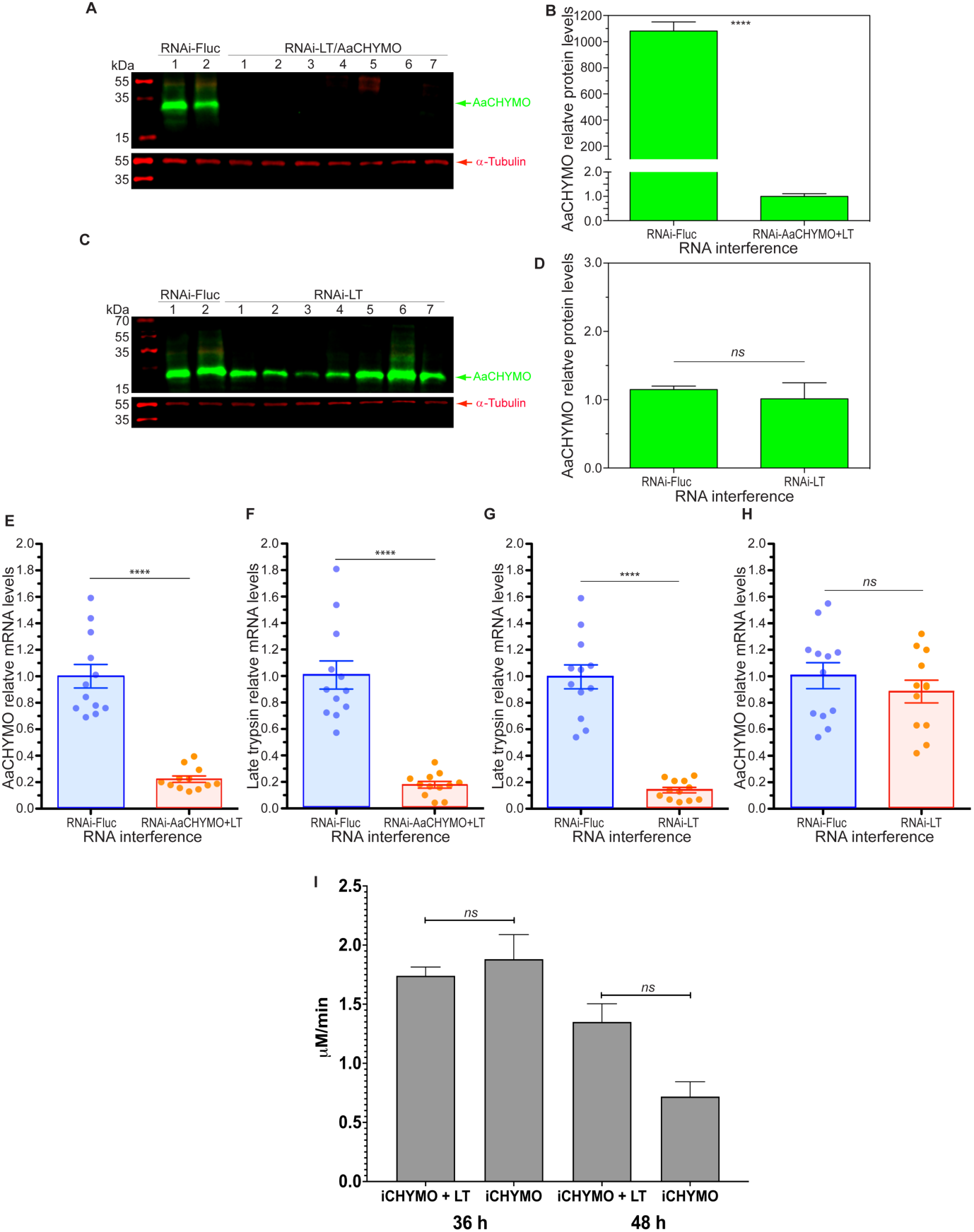
Double RNAi knockdown of AaCHYMO and AaLT in individual *Ae. aegypti* mosquitoes. **A.** Western blot analysis showing that the double knockdown led to an effect on AaCHYMO expression (seven individual mosquitoes) and compared to the FLUC control (two individual mosquitoes). **B.** Quantification of AaCHYMO protein expression levels in midgut extracts with the RNAi double knockdown of AaCHYMO and AaLT. **C.** Western blot analysis showing that knockdown of AaLT has no effect on AaCHYMO expression. **D.** Quantification of AaCHYMO mRNA levels in mosquito midgut tissues with the RNAi double knockdown of AaCHYMO and AaLT. **E.** Quantification of AaLT mRNA levels in mosquito midgut tissues with the RNAi double knockdown of AaCHYMO and AaLT. **F.** Quantification of AaLT mRNA levels in mosquito midgut tissues with the RNAi knockdown of AaLT only. **G.** Quantification of AaCHYMO mRNA levels in mosquito midgut tissues with the RNAi knockdown of AaLT only. Experiments in **B**, **D**, and **E-G** were conducted from 3 biological cohorts using 5 to 12 mosquitoes. Data are presented as mean ± SEM. Statistical significance is represented by stars above each comparison (unpaired Student’s t-test; **** p < 0.0001 compared to the RNAi-FLUC control samples, ns = no significance). **I.** Activity assays comparing the RNAi AaCHYMO alone knockdown versus the AaCHYMO/AaLT double knockdown using the MeO-Suc-Arg-Pro-Tyr-AMC substrate. No significant changes in overall activity were observed in the double knockdown (4 to 7 biological cohorts using 3 mosquitoes were tested).

Given the possibility that individual knockdown of midgut proteases can lead to a direct effect on fecundity ^1^, maximal egg production in individual RNAi AaCHYMO mosquitoes was measured. The absence of AaCHYMO did not affect egg production (**Supplementary Figure 2**).

### Human blood protein digestion assays reveal AaCHYMO cannot process globular proteins

Further characterizing the role of AaCHYMO in blood meal protein digestion, we co-incubated the enzyme with HSA, Hb, and IgG. These blood proteins are known to be processed into polypeptides and amino acids used by the mosquito ^1^. To detect potential cleavage products by AaCHYMO a 10:1 (protein to protease) ratio was set and compared to the activity of human and bovine chymotrypsin. Interestingly, AaCHYMO did not readily cleave all three blood meal proteins like human and bovine chymotrypsin (**Figure 4**). For instance, in the HSA reactions, AaCHYMO digestion led to only a single cleavage product running near 30 kDa (teal arrow in **Figure 4A**) and no notable decrease in overall intact HSA. Conversely, multiple cleavage products were observed for human and bovine chymotrypsin, as well as a visible decrease in the amount of intact HSA. Similarly, a single cleavage product by AaCHYMO was observed near 30 kDa (red arrow in **Figure 4B**) without digesting any more Hb. Human and bovine chymotrypsin cleaved Hb leading to several cleavage products observed below 15 kDa (**Figure 4B**). The mammalian chymotrypsin enzymes also cleaved the Hb species above 30 kDa (blue arrow in **Figure 4B**). Lastly, in the IgG reactions (**Figure 4C**), various cleavage products were observed for human and bovine chymotrypsin, but only one single cleavage product observed at 15 kDa for AaCHYMO (aqua arrow in **Figure 4C**). Two additional cleavage products were observed between 50 and 70 kDa in all three reactions (red and orange arrows in **Figure 4C** as a representative example), which were readily digested by human and bovine chymotrypsin.

**Figure 4.**
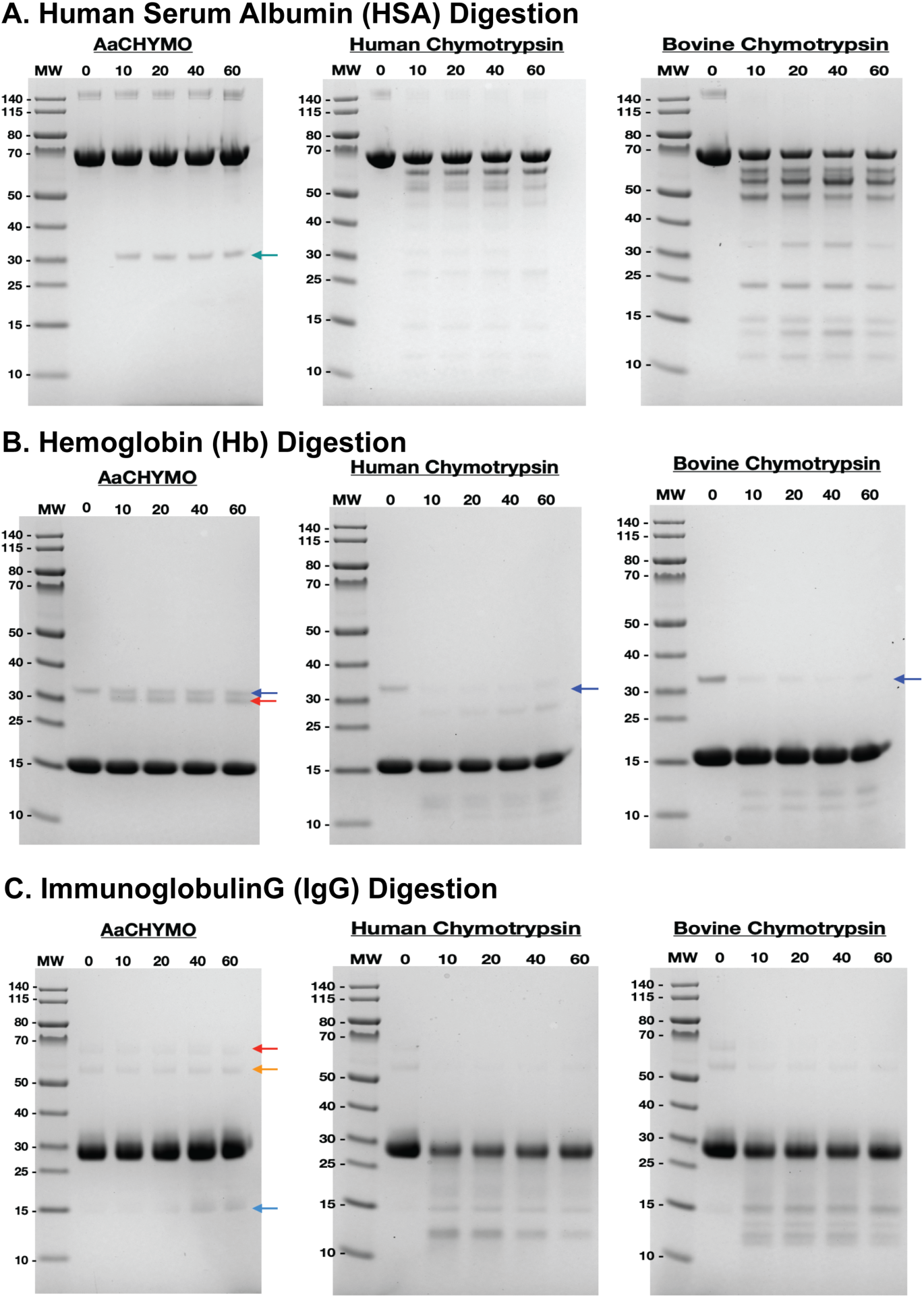
Protein gel electrophoresis analysis of AaCHYMO, human and bovine chymotrypsin digestion assays with the three major human blood proteins: human serum albumin (HSA) (**A**), hemoglobin (Hb) (**B**), and immunoglobulin G (IgG) (**C**), incubated for a total of 60 min post-protease addition. Time = 0 min is the no protease control in each reaction. **A.** Comparing the HSA digestion fragments of AaCHYMO only one single cleavage product is observed (teal arrow), as opposed to several digestion fragments observed in human and bovine chymotrypsin digestion assays. **B.** In the Hb digestion assays, only one single cleavage product is observed (red arrow), while the higher molecular weight (MW) Hb band protein species (blue arrows) stays intact throughout the whole reaction time but is readily degraded by human and bovine chymotrypsin. **C.** In the IgG digestion assays, the two top bands in between the 50 and 70 kDa MW marker (red and orange arrows) remain intact throughout the reaction time, with only a single cleavage product observed appearing at 20 min after protease addition (aqua arrow). Human and bovine chymotrypsin readily degrade the two bands between the 50 and 70 kDa MW marker (red and orange arrows in the AaCHYMO digestion reaction), with more observable cleavage products below the 30 kDa MW marker observed.

With these surprising digestion results, we decided to analyze the midgut protein content in the RNAi AaCHYMO- and FLUC-injected mosquito midgut tissue extracts used for the detection of proteolytic activity (**Figures 1** and **2**). Analyzing this content should reveal differences in the digestion of blood proteins if AaCHYMO is absent. Only three selected protein species found in all gel samples were analyzed and are shown in representative gels for both FLUC and RNAi AaCHYMO (see purple, red, and teal arrows in **Figure 5A**). Overall, there are no significant differences in the digestion of the three proteins (**Figures 5B-5D**). In fact, all gels (FLUC *vs* RNAi AaCHYMO) did not display notable differences, except for the 3 h timepoint of RNAi AaCHYMO *vs* FLUC (p value < 0.05) with protein products found between 50 and 70 kDa (red arrow in **Figure 5A** and data in **Figure 5C**), where the band in the RNAi AaCHYMO is visibly lighter than the other timepoints.

**Figure 5.**
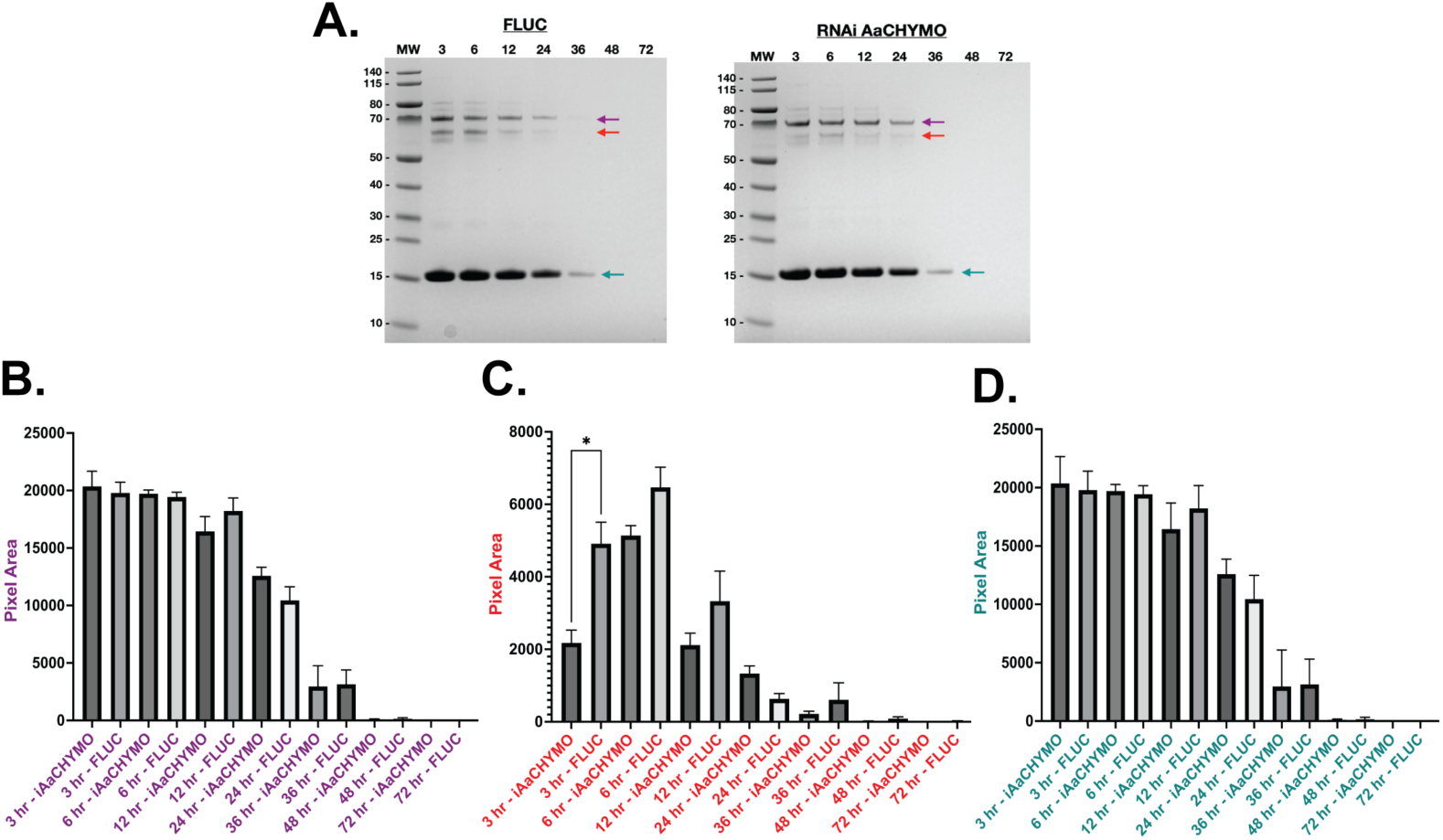
*Ae. aegypti* midgut blood protein digestion extract analysis. **A.** Representative protein gels of FLUC and RNAi AaCHYMO midgut extracts collected at 3 h to 72 h PBF (0.006 midgut equivalents were loaded onto a NuPAGE 4-12% BIS-TRIS gel (Invitrogen)). MW represents the pre-stained PageRuler protein ladder in kilodaltons (kDa) (Thermo Scientific #26616). **B.** Column plot comparing the pixel area of the protein species running at the 70 kDa MW marker (purple arrow in Figure 5A) between RNAi AaCHYMO and FLUC midgut extract samples. **C.** Column plot comparing the pixel area of the protein species running between the 50 and 70 kDa MW marker (red arrow in Figure 5A) between RNAi AaCHYMO and FLUC midgut extract samples. **D.** Column plot comparing the pixel area of the protein species running at the 15 kDa MW marker (teal arrow in Figure 5A) between RNAi AaCHYMO and FLUC midgut extract samples.

### Human blood protein digestion specificity is supported by molecular docking simulations

Aiming to predict the potential interactions of AaCHYMO with the three suggested substrates (HSA, Hb, and IgG), we employed a consensus with three different docking software. First, docking simulations using HDOCK predicted a few potential interactions for HSA with AaCHYMO (**Figure 6A**), which were not within feasible distances for possible interactions (∼5 Å) ^19,20^ close to the catalytic site. A docking score of −193.74 and a confidence score of 0.70 were predicted, which could suggest HSA is likely to interact, as protein-protein complexes in the PDB have a docking score of around −200 and are likely to interact when having a confidence score above 0.7 ^21^. Similarly, Hb was predicted likely to interact, having a score of −208.6 and confidence of 0.76, but no feasible interactions were observed close to the catalytic serine (**Figure 6B**). These results suggest that the two human proteins can potentially engage, but inconclusive for full cleavage to occur. Conversely, IgG is predicted to likely interact (−252.11 and 0.88) with AaCHYMO (**Figure 6C**), especially at the edge of both its light (−284.26 and 0.93) and heavy chains (−319.24 and 0.96) and being closer to the catalytic site (∼7 Å, ranging from 5.3 to 9 Å). Also, additional interactions are predicted around the catalytic site, which could support the stabilization of these contacts.

**Figure 6.**
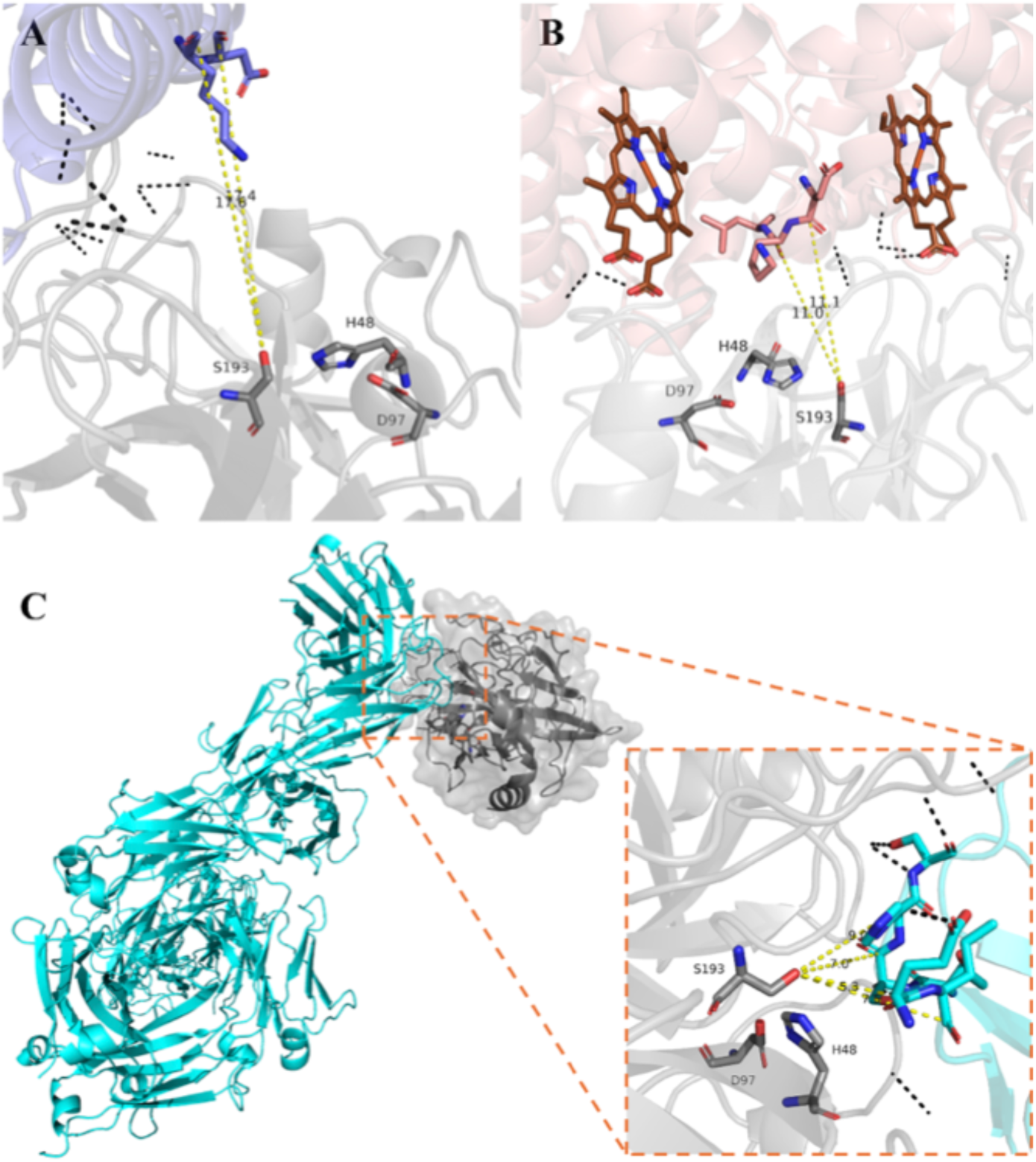
AaCHYMO is predicted to interact with IgG and less likely to interact with human albumin or hemoglobin. Modeled structure of AaCHYMO by AlphaFold2 (gray) was submitted to HDOCK to predict potential interactions with (**A**) human serum albumin (PDB ID: 1E7H, slate blue), (**B**) human hemoglobin (PDB ID: 1GZX, salmon), and (**C**) human neutralizing IgG (PDB ID: 1HZH, cyan). Predictions suggest the latter’s potential to interact and have residues close to the catalytic site of the enzyme (∼7Å). Results suggest that the two other human proteins can potentially engage, but inconclusive for full cleavage to occur. Predicted interactions are shown as black dashed lines, while measured distances to the catalytic serine are shown as yellow dashed lines. Heme groups are displayed as sticks (brown). Catalytic residues are displayed as sticks (gray) and labeled (H48, D97, and S193). Structures are represented by cartoons and a transparent surface. All images were generated with PyMOL software (v2.5.7).

Consistently, simulations using HADDOCK 2.4 were able to find 15 potential structures for IgG interactions with AaCHYMO, grouped in seven different clusters suggesting their potential PPI, while only four potential structures in two clusters were found for both HSA and Hb. Notable score differences between each cluster, which are supportive of PPI ^22^, were observed for IgG (ranging from −55.2 ± 7.2 to −16.4 ± 0.7) and albumin (−42.1 ± 8.8 and −28.7 ± 5.9), but not for hemoglobin (−57.7 ± 2.3 and −50.2 ± 5.5). Lastly, the three proteins are not predicted to interact with AaCHYMO by PEPPI simulations. Herein, HSA, Hb, and IgG structures were predicted to have negative likelihood ratios (LR < 0) of −2.412, −2.331, and −2.045, respectively, which usually classify two proteins as non-interacting ^23^. Interestingly, however, IgG had the highest difference (9.192 *vs.* 1.554) between the positive prediction score (structure similarity, *i.e.*, SPRING) *vs.* the negative prediction score (non-interaction similarity, *i.e.*, SPRINGNEG)^23^. This could suggest a higher potential of IgG to interact with AaCHYMO when compared to albumin (3.673 *vs.* 2.212) and hemoglobin (4.782 *vs.* 3.376). Overall, these docking simulations for PPI support our digestion assays (**Figure 4**).

### Knockdown of EcR reveals possible regulation of AaCHYMO expression and activity

To explore the potential role of EcR in AaCHYMO protease regulation, knockdown studies were conducted and chymotrypsin-like activity detected using the MeO-Suc-Arg-Pro-Tyr-AMC substrate. The knockdown of EcR led to an overall decrease in AaCHYMO protease expression when compared to the FLUC-injected control (**Figure 7A**-**7C**), and surprisingly, a significant reduction in chymotrypsin-like activity in midgut tissue extracts at 6 h, 24 h, and 36 h PBF (**Figure 7D**).

**Figure 7.**
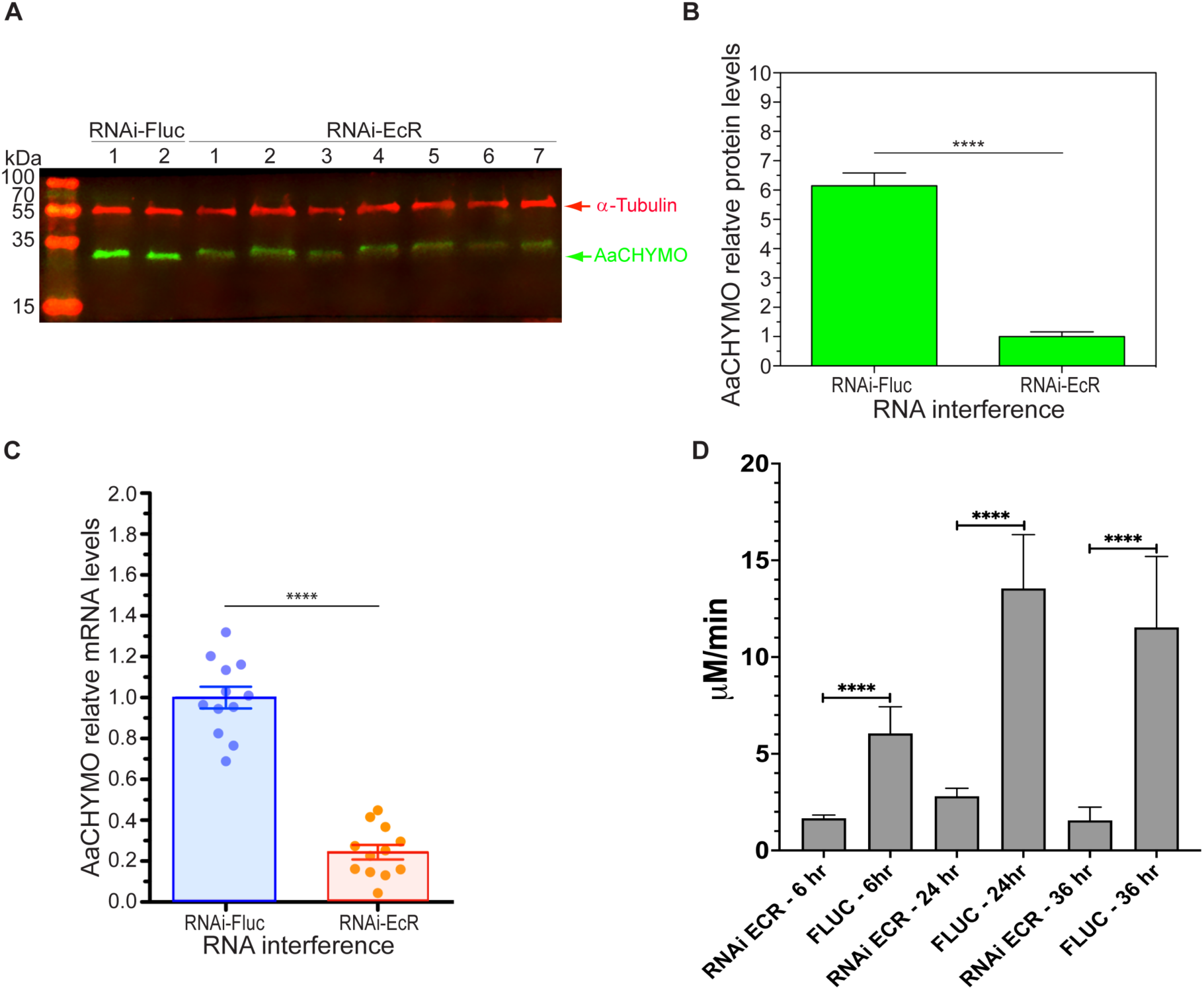
RNAi ecdysone receptor (EcR) knockdown effects on AaCHYMO protease expression and activity. **A.** Western blot analysis of the EcR knockdown effect on AaCHYMO. **B.** AaCHYMO protein expression levels were quantified using ImageJ and are shown to be less than the FLUC control samples. **C.** Quantification of AaCHYMO mRNA levels in EcR knockdown mosquito tissue extracts compared to the FLUC control. Experiments in **B** and **C** were conducted from 3 biological cohorts using 5 to 12 mosquitoes. Data are presented as mean ± SEM. Statistical significance is represented by stars above each comparison (unpaired Student’s t-test; **** p < 0.0001 compared to the RNAi-FLUC control samples). **D.** Chymotrypsin-like activity comparison between RNAi EcR and the FLUC control midgut extract samples using the MeO-Suc-Arg-Pro-Tyr-AMC substrate. Significant reduction (****p < 0.0001) in overall activity is observed in the EcR knockdown samples (3 to 4 biological cohorts using 3 mosquitoes were tested).

## DISCUSSION

The most studied endopeptidases and major contributors in blood meal protein digestion in the *Ae. aegypti* mosquito midgut are trypsin-like enzymes, with Arg and Lys amino acid preferences at the P1 site ^1,3–5,24^. Other endopeptidases found in the midgut are serine collagenase-like ^2–4^ and chymotrypsin-like enzymes ^6–8^, but their overall contribution to blood protein digestion is still relatively unknown. Here, we aimed to understand the complex nature of proteolytic blood protein degradation by biochemically studying AaCHYMO *in vitro* and *in vivo*, the first chymotrypsin-like enzyme identified in 1997 ^6^. This is the first step in validating if all or a select few midgut proteolytic enzymes could serve as potential inhibitor targets for the development of a new vector control strategy.

With the success of solubly expressing, activating, and purifying AaCHYMO, activity was tested using several commercially available chymotrypsin substrates. Obvious specificity differences were observed between all three (including human and bovine) chymotrypsin-like enzymes. The most preferred amino acids at the P1 position for human and bovine chymotrypsin are Phe, Trp, and Tyr, with lower preference for Leu and Met ^25,26^. Similarly, AaCHYMO readily cleaved substrates with Phe, Tyr, Trp, Leu, and Met at P1, and only human chymotrypsin cleaved Arg and Lys substrates. This is not surprising given that the protease has been shown to cleave peptide substrates with an Arg or Lys at P1 but with an overall lower rate than when aromatic amino acids are found at this position ^27^. Additionally, depending on which amino acids are present at P2-P4, the specificity and activity towards the preferred amino acids at P1 can be influenced ^27,28^. For example, in bovine chymotrypsin, when comparing Suc-Ala-Ala-Pro-Phe-AMC with Suc-Ala-Ala-Phe-AMC, the specificity constant (*k_cat_/K_m_*) is nearly 21-fold higher when Pro is added at P2. Similar results for human chymotrypsin have been observed ^27,28^. As for AaCHYMO, the preferred order of amino acids at P1 was determined to be Tyr, Leu, Phe, and Met using MSP-MS ^3^, which match the *in vitro* activity assays using commercially available chymotrypsin substrates in this study.

Moreover, like human and bovine chymotrypsin, certain amino acids at P2-P4 influence the activity and specificity at P1 in AaCHYMO. For instance, when comparing the specificity constants (*k_cat_/K_m_*) of AaCHYMO with commercially available substrates to human and bovine chymotrypsin, *k_cat_/K_m_* is overall lower than both mammalian chymotrypsin enzymes (see the ratio comparisons in **Table 1**). Thus, we synthesized AaCHYMO-specific preferences at the P2-P4 sites and one substrate with a non-preferred amino acid at P2 (Ac-Ala-His-Asp-Trp-AMC). When the AaCHYMO-specific substrates were used, in most cases, the *k_cat_/K_m_* ratio changed to favor AaCHYMO, with substrates having Leu and Met at P1 being processed more readily than human and bovine chymotrypsin. In addition, when comparing the *k_cat_/K_m_* values of the AaCHYMO-specific substrate reactions, the values were greater than those determined using the commercially available substrates. As for the non-preferred amino acid substrate at P2 (Asp in this case), AaCHYMO was not able to cleave at the P1 position even though a highly preferred Trp is found at P1. Both human and bovine chymotrypsin readily and favorably cleaved the Ac-Ala-His-Asp-Trp-AMC leading to *k_cat_*/*K_m_* greater than 3×10^6^ M^-1^ s^-1^. These results indicate the importance of knowing the amino acid preferences beyond the P1 position (P2-P4), especially if specific protease inhibitors are to be designed to target individual midgut proteases. Lastly, by performing these *in vitro* activity assays, we have confirmed that AaCHYMO is indeed a chymotrypsin that cannot cleave peptides with Arg or Lys at P1.

Furthermore, we set to test the overall chymotrypsin-like activity of midgut tissue extracts from sugar- and blood-fed mosquitoes, as well as in pupae, adult male mosquitoes, female carcasses (without the midgut), and midgut tissues without the food bolus (-FB). Activity was detected in the 3 h to 48 h PBF midgut tissue extracts, which corresponds to the presence of the active form of AaCHYMO. These results corroborate the work of Jiang *et al*. ^6^, who first identified the mRNA and protein expression of AaCHYMO in blood fed mosquito extracts. AaCHYMO mRNA is only present in 24 h post-emerged female mosquitoes with the protease being translated only after ingestion of blood ^6^. Interestingly, the Jiang *et al.* study did not detect the presence of AaCHYMO in sugar-fed (or unfed) mosquitoes, and in our case, AaCHYMO was clearly present in these samples, but as the inactive zymogen form, with no detectable chymotrypsin-like activity. However, chymotrypsin-like activity increased after ingestion of a blood meal, with maximal activity at the 12 h to 36 h timepoints. As for mosquito tissues tested at different life stages, very low negligible activity was observed in pupae, adult males, female carcasses (without the midgut), and the midgut without the food bolus. However, in larvae and the food bolus, higher overall activity was observed. In fact, the food bolus has higher activity than even the midgut extract tissue samples from **Figure 1D**. These results are not surprising since active AaCHYMO is strictly expressed for release into the midgut after ingestion of a blood meal ^6^. The food bolus sample is expected to show such high activity as it contains all secreted proteases released to help digest blood meal proteins, and may include activity from JHA15, AaCHYMO II, AaLT, and other midgut proteases with unknown chymotrypsin-like activity. The activity detected in the larval stages may correspond to AaCHYMO II, which was shown to be strictly found in both larval and adult stages of *Ae. aegypti* ^8^. However, future studies are needed to confirm the activity of AaCHYMO II in the midgut.

Knockdown studies of AaCHYMO did not lead to an effect on fecundity, but did lead to the complete elimination of chymotrypsin-like activity in the 3 h to 24 h PBF samples, and a significant decrease in activity at 36 h PBF (p < 0.0001) and 48 h PBF (p < 0.05). It is important to note that many midgut proteases are expressed and released in the mosquito midgut at the same time as active AaCHYMO. In particular, AaLT is expressed in the late phase starting at 18 h PBF and present until 36 h PBF ^1^. With this knowledge, we performed a double knockdown of AaLT and AaCHYMO. Surprisingly, this did not lead to a change in overall chymotrypsin-like activity, indicating that a different protease or a different set of proteases may be giving rise to the residual activity. The other possibilities are JHA15 ^7^ and AaCHYMO II ^8^, the two additional chymotrypsin-like enzymes expressed in the mosquito midgut. Proteolytic profiling and the study of these two proteases should shed light on their role in blood meal digestion and activity at the 36 h and 48 h PBF time points.

AaCHYMO was incubated with HSA, Hb, and IgG, and the activity compared to human and bovine chymotrypsin. To our surprise, AaCHYMO led to the release of only one cleavage product in each reaction, but even then, the fragment from each reaction was not further processed by the protease. In contrast, human and bovine chymotrypsin digested HSA, Hb, and IgG, releasing several cleavage products in each reaction. This may suggest AaCHYMO is highly selective or it requires cooperation from other proteases to expose certain amino acids for complete blood meal protein cleavage. To understand if potential PPI would occur between AaCHYMO and the different blood proteins we predicted their interactions using different docking simulations in a consensus approach. The HDOCK predictions with the three human blood proteins revealed IgG being the only protein likely to interact with AaCHYMO, which was also supported by simulations using HADDOCK 2.4 and less so by PEPPI. Our simulations supported the enzyme cleavage observed from our digestion assays, where we identified the release of an IgG cleavage product over time (see **Figure 4C**). Conversely, HADDOCK 2.4 and PEPPI both predicted that Hb was not likely to interact with AaCHYMO. Despite being likely to interact based on HDOCK predictions, Hb residues were not within feasible distances close to the catalytic serine, which interestingly, in the AaCHYMO/Hb digestion assay, a single cleavage product was observed (see **Figure 4B**). Lastly, despite one product being formed after cleavage of HSA in our digestion assays, the simulations were inconclusive to the likeness of albumin to interact with AaCHYMO, having few potential interactions predicted and less likely to occur. One could argue that the confirmational changes of albumin in solution at various pH ^29,30^ could support its interaction with AaCHYMO and subsequent cleavage to occur. Arguably, this could explain our prediction outcomes, which considers only the available crystallographic data ^31^ for the simulations.

The lack of fecundity effects by the absence of AaCHYMO and the lack of individual blood protein digestion by the protease can be explained by the presence of other midgut proteases that may compensate and work collectively to degrade blood proteins. For instance, in nematode, trematode, and plasmodia parasitic organisms, blood protein digestion follows a protease network (or cascade) involving several proteases (for an extensive review see ^32^). In *Ancylostoma caninum* and *Plasmodium falciparum*, aspartic and cysteine proteases must first make initial cuts on hemoglobin, which is then further processed by metalloproteases to fully digest the blood protein ^32^In *Ae. aegypti*, after ingestion of a blood meal, several trypsin-like proteases (AaET, AaSPII-AaSPIV), chymotrypsin-like proteases (AaCHYMO, JHA15, AaCHYMO II), serine collagenase-like protease (AaSPI, AaSPV), as well as exopeptidases (amino- and carboxypeptidases) are expressed and released in the early phase (0 h to 12 h PBF), allowing these proteases to work collectively and start digesting globular blood proteins ^1,2,4,8,17,33^. The mosquito does not complete digestion until after the second (or late) phase (approximately 40 h PBF) where active AaCHYMO is still present, along with a new set of trypsin-like proteases (AaSPVI and AaSPVII), the constitutively expressed AaSPII-AaSPV proteases, continued expression of exopeptidases, and AaLT ^1–3,17^. Without the collective activity of early and late phase midgut proteases, blood protein digestion would not be effective and efficient, and the mosquito would not have the required nutrients to fuel the gonotrophic cycle. So, it is not surprising that when AaCHYMO was knocked down, there was no overall effect on digestion or fecundity. The proteolytic network in the mosquito midgut compensated for the absence of the protease, allowing the mosquito to fully digest the blood proteins needed for nutrition. On the other hand, knockdown of three late phase midgut proteases (AaSPVI, AaSPVII, and AaLT ^1^) did have a direct effect on fecundity, therefore, it is important to study all proteases involved to delineate which would be effective targets for the development of inhibitors.

As alluded to above, the ingestion of blood by the female mosquito leads to broad yet crucial physiological consequences ^34,35^. As blood meals are strictly linked to mosquito development ^36,37^ and reproduction ^1,38^, there is tight coordination between large cohorts of genes and mosquito biology in a spatiotemporal manner. Blood meal digestion occurs in a biphasic manner, and designated midgut proteases are expressed and active to digest blood proteins. Early phase protease expression is proposed to be translationally regulated while late phase protease expression is transcriptionally regulated ^39^. However, the exact mechanisms or the players involved are still relatively unknown. The 20-hydroxyecdysone (20-E) master regulator and its interacting partner EcR are known to be involved in fecundity and oviposition ^18^. Expression of the nuclear receptor EcR is found to increase 1-6 h PBM and again after 24 h PBM in the fat bodies and ovaries of *Ae.* aegypti female mosquitoes ^40^, which correlates with midgut protease digestion of blood proteins ^1^. Furthermore, RNAi knockdown of midgut EcR significantly decreased mRNA transcript levels of *SPI*, *SPVI*, *SPVII*, and *LT* midgut proteases ^17^. AaCHYMO was not included in the study, and thus, we performed RNAi knockdown on EcR to observe its effect on AaCHYMO expression and activity. The knockdown of EcR led to lower AaCHYMO protein expression and significantly reduced chymotrypsin-like activity, therefore illustrating EcR’s possible role in regulating not only AaCHYMO expression and activity, but its effect on other midgut proteases. More studies focusing on EcR are needed to delineate the true role in regulating AaCHYMO and overall midgut protease expression.

Blood protein digestion in *Ae. aegypti* was thought to follow a simple biphasic release of midgut proteolytic enzymes, with some proteases translated almost immediately after a blood meal in the early phase (translationally controlled), followed by transcription and translation of the next set in the second phase (transcriptionally controlled) ^1,2^, with most of the activity coming from trypsin-like proteolytic enzymes ^1–5^. However, with the discovery of other serine proteases ^2,7,8^, the mosquito seems to have evolved a complex network of proteolytic enzymes, each having a broad range of substrate specificities and activity profiles to efficiently process blood within 40 h PBF. If one is to target midgut proteolytic enzymes as potential inhibitor targets, then it is crucial to elucidate the activity profile and role of all midgut proteases. As was shown in this study, AaCHYMO is a true chymotrypsin-like enzyme cleaving Phe, Tyr, Trp, Leu, and Met at P1, but not Arg or Lys. Furthermore, knockdown of the protease did not influence fecundity or affect blood protein digestion, even though chymotrypsin-like activity was eliminated and significantly reduced. These results indicate the importance of carefully elucidating the complex network of midgut proteases that compensate for the absence of AaCHYMO. Given that no phenotypic changes were observed with the knockdown, it is likely that the mosquito evolved a redundancy in function ^7^ to ensure complete blood meal digestion. In addition, regulation of protease expression and activity by EcR must be fully elucidated to determine if this nuclear receptor not only affects AaCHYMO but other understudied proteases. Therefore, it is important to also characterize the activity profiles and roles of AaSPI-AaSPV ^2^, JHA15 ^7^, and AaCHYMO II ^8^, their mode of regulation, and effect on fecundity.

## METHODS

### Chemicals and Fluorescent Substrates

TRIS base (J.T. Baker #JT4099-6), Calcium chloride (CaCl_2_) (J.T. Baker # JT1332-1), Tween 20 (MilliporeSigma #80503-492), 7-amino-4-methylcoumarin (AMC) (Thermo Scientific #AAA15017MD), PageRuler Prestained Protein Ladder (Thermo Scientific #PI-26616). **Supplementary Table 1** lists all commercially available chymotrypsin and the AaCHYMO-specifically designed fluorescent AMC substrates used in this study. AaCHYMO-specific substrates were synthesized by Biomatik USA, LLC (Wilmington, DE) based on the identification of preferred amino acids at the P1 to P4 sites ^3^.

### AaCHYMO Enzymatic Assays and Kinetic Parameter Determination

Recombinant AaCHYMO was expressed in SHuffle T7 Express Competent *Escherichia coli* Cells (New England Biolabs #C3029J), purified using a 5 mL HisTRAP FF column (Cytiva #17525501) on the AKTA Pure 25 L (Cytiva, Marlborough, MA), and activated via buffer exchange, as described ^3^. Kinetic parameters were then determined in activity assay buffer containing 20 mM TRIS-HCl pH 7.2 and 0.1% Tween 20 (30 μL total volume in a 384-well black microplate (Thermo Scientific # 262260)). Active AaCHYMO concentrations were in the range of 5 to 100 nM and incubated with varying commercially available chymotrypsin fluorescent substrate concentrations or specific-AaCHYMO designed fluorescent substrates (**Table 1**) (under steady-state conditions). For comparison, 50 pM to 50 nM human pancreas chymotrypsin (Athens # 16-19-030820) and bovine chymotrypsin alpha (Abnova # P5212) were tested under the same reaction conditions. Reaction rate slopes were measured in triplicate using the Tecan Spark (Männedorf, Switzerland) at 360 nm excitation and 460 nm emission wavelengths. Chymotrypsin activity was monitored by the release of 7-amino-4-methylcoumarin (AMC). A standard curve using AMC was used to convert relative fluorescence units per sec (RFU’s/sec) to a change in product concentration over time (μM/min). Briefly, AMC stocks from 0 to 100 μM (serial dilution) were prepared in activity assay buffer and the fluorescence at each concentration was measured at 360 nm/460 nm in duplicates. The values were plotted on Excel and the slope determined using a linear regression plot. To compare all chymotrypsin data, a final 5 nM enzyme concentration was used to estimate kinetic constants (*v_max_*, *k_cat_*, and *K_m_*) using Michaelis-Menten non-linear regression analysis at a 95% Confidence Interval (CI) on GraphPad Prism version 10.4.0. Comparison assays between AaCHYMO, human and bovine chymotrypsin, were performed using substrates that were specifically designed for AaCHYMO. The lower level of AMC detection was determined by preparing a serial dilution from 1 nM to 1 pM AMC in activity assay buffer, as recommended in the Tecan Spark manual. Dilution fluorescent readings at each concentration was measured at 360 nm/460 nm in duplicates. The values were plotted on Excel and the slope determined using a linear regression plot.

### Mosquito Rearing, Blood feeding, and Tissue Preparation

*Ae. aegypti* mosquitoes (Rockefeller strain) were maintained on a 10% sucrose diet until feeding, with rearing conditions set at room temperature (25°C), 80% relative humidity, and a 16 h light: 8 h dark cycle. Whole bovine blood with citrate anticoagulant (HemoStat #BBC500, Dixon, CA) was used to blood feed mosquitoes. Mosquito midguts were dissected in 1x PBS under a light microscope and transferred to 20 mM TRIS-HCl pH 7.2 buffer. Three midguts per 100 μL was prepared for a final concentration of 0.03 midgut/μL. Mosquito midgut tissues were dissected at 3-, 6-, 12-, 24-, 36-, 48-, and 72-hours post-blood feeding (PBF), and as a control, sugar fed midgut extracts were used. Other samples isolated were the *Ae. aegypti* female carcass (without the midgut and food bolus), midgut minus the food bolus, the food bolus only, larvae, pupae, and adult male mosquitoes. Three of each of these samples were also processed as the midgut extracts, 3 per 100 μL of 20 mM TRIS-HCl pH 7.2. All samples were immediately flash frozen in liquid nitrogen and stored at −80°C until needed.

### Synthesis of double-stranded RNA (dsRNA)

Genes encoding *Ae. aegypti* AaCHYMO (EAT45655; AAEL003060), AaLT (EAT34478; AAEL013284), and EcR (EAT38529; AAEL009600) were identified from the mosquito genome in the NCBI database. The T7 promoter sequence (5’ TAATACGACTCACTATAGGGAGA 3’) was added to the 5’-end of both the forward and reverse primers for each gene **(Supplementary Table 2)**. PCR was performed to amplify the dsRNA region of the DNA and sequences verified before use. dsRNA targeting each gene were synthesized using the NEB HiScribe T7 RNA Synthesis kit (New England Biolabs #E2040S) with PCR amplified DNA templates. Purified dsRNA was then resuspended in nuclease-free HPLC water at 7.5 μg/μL and stored in the −80°C freezer until use. *Ae. aegypti* females were microinjected with 2.0 μg of dsRNA using a Nanoject II micro-injector (Drummond Scientific Company, Broomall). The injection was performed within 4 hours after adult eclosion, and the dsRNA microinjected mosquitoes were maintained on 10% sucrose until needed for experiments. AaCHYMO, LT, a combination of AaCHYMO/LT, and EcR dsRNA microinjections were performed. The firefly luciferase dsRNA microinjection was used as a negative control.

### Processing of Mosquito Tissues and Chymotrypsin-like Activity Determination

All isolated wildtype (WT) and dsRNA injected mosquito tissues were processed via sonication utilizing the Sonic Dismembrator (Fisherbrand Model 120) with a 5/64-inch microtip. Samples were thawed on ice, followed by quick vortex mixing (10 sec) and centrifugation (using the mini-table top centrifuge for 10 sec), then samples lysed using the dismembrator (25% amplitude (AMP)) for 10 sec (3x total). To avoid overheating, samples were set on ice for about 30 sec, while the other samples were being processed. The microtip was cleaned with ultrapure water and wiped dry with a Kimwipe tissue each time before the next sample was processed to avoid cross-contamination. For the female carcass and the adult males, samples were sonicated 5 times to ensure proper lysis. After lysing, all samples were then centrifuged at full speed in the VWR MicroStar 21 temperature-controlled centrifuge (14,800 rpm, 8°C) to remove any tissue, particles, or contaminants away from the lysate. The lysates were then transferred to a new tube and flash frozen in liquid nitrogen and stored at −80°C until needed.

To test activity in all WT and dsRNA-injected mosquito tissue extracts, the best commercially available substrate processed by AaCHYMO (MeO-Suc-Arg-Pro-Tyr-AMC) was used (see **Table 1**). Given that mosquito midgut extracts contain several midgut proteases that can potentially process the substrate, either a 250- or 10,000-fold dilution in activity assay buffer (20 mM TRIS-HCl pH 7.2 and 0.1% Tween 20) was prepared to test for activity (30 μL reactions, similar setup as above for the recombinant AaCHYMO kinetic parameter assays). However, the final substrate concentration used was 25 μM to ensure linear regression activity during the run. A minimum of 3-6 replicates were performed for nearly all tissue extracts. Data was statistically analyzed by unpaired Student’s t-test with mean values ± SEM using GraphPad Prism version 10.4.0. Asterisks indicate significant differences (**p < 0.05; ****p < 0.0001; ns, not significant).

### Western Blot Analysis of Mosquito Midgut Tissue Extracts

A polyclonal antibody against AaCHYMO was prepared by GenScript (Piscataway, NJ) using the antigenic AaCHYMO protein sequence without the signal/leader peptide. Western blotting was performed as previously described for other mosquito proteins ^41,42^. Briefly, midgut protein extracts (midgut epithelial cells without the food bolus) were separated via 12% SDS-PAGE gels, electroblotted onto nitrocellulose membranes (LI-COR Biosciences), incubated with AaCHYMO and α-tubulin antibodies at 4°C overnight. The membranes were washed with 1x PBST and incubated with the secondary antibodies for 1 hour at room temperature. Visual detection was performed with the Odyssey XF Infrared Imaging System (LI-COR Biosciences). The dilutions of the primary antibodies were as follows: AaCHYMO antibody (1:1000) and α-tubulin antibody (1:3000). The secondary antibodies (LiCor IRDye 800CW goat anti-rabbit and LiCor IRDye 680RD donkey anti-mouse) were diluted at 1:10000. ImageJ software was used to quantify protein band intensity. Each band intensity was normalized to α-tubulin band signals. Experiments were conducted from 3 biological cohorts using 5 to 10 mosquitoes. For AaCHYMO time course studies, the protein levels were analyzed by ANOVA with Tukey’s post-hoc test. Unpaired Student’s *t*-test using GraphPad Prism was used to determine whether there is a statistical difference between the two RNAi groups.

A chemiluminescent WB analysis of midgut extracts (midgut epithelial cells with the food bolus), which are the representative samples used for the detection of chymotrypsin-like activity, was also done. Midgut samples were sonicated and prepared as describe above. For this WB, 0.5 midgut equivalents were used, separated using a 4-12% BIS-TRIS gel (12-well, Invitrogen # NP0322BOX), electroblotted to a PVDF membrane (Millipore Sigma Immobilon-PSQ #ISEQ00010), incubated with a 1:1500 dilution of AaCHYMO primary antibody at 4°C overnight. The membrane was washed with 1x TBST and incubated with a 1:2000 dilution of the secondary antibody (Amersham ECL Anti-Rabbit IgG #NA934-1ML) for 1 hour at room temperature. Visual detection was performed with the chemiluminescent setting on the BioRad ChemiDoc MP Imaging System (Hercules, CA) after incubating the membrane with the SuperSignal West Pico Chemiluminescent Substrate (Thermo Scientific #PI34577) for 30 sec.

### Validating RNAi Knockdown Efficiency

Single mosquito analysis was used to verify knockdown efficiency from dsRNA-microinjected mosquitoes. Midgut tissue samples were dissected after blood feeding. Proteins were extracted in 1x lysis buffer, while total RNAs were isolated with TRIzol (Invitrogen). Optical densities of total RNAs were determined using the Nanodrop (Thermo Scientific). DNaseI-treated total RNAs (200 ng) were used to synthesize complementary DNA (cDNA) using an oligo-(dT)-VN primer and reverse transcriptase (Promega). Quantitative real-time PCR (qPCR) was performed with Maxima SYBR Green qPCR Master Mix (Thermo Scientific) and gene-specific primers (final concentration at 200 nM, **Supplementary Table 2**) using the CFX Connect qPCR instrument (BioRad). Knockdown efficiency was compared using FLUC dsRNA-microinjected female control mosquitoes. Relative mRNA expression for AaCHYMO, EcR, and LT were normalized to ribosomal protein S7 mRNA levels in the same cDNA samples. Data was obtained from individual midgut of 12 mosquitoes. Data are presented as mean ± SEM. Statistical significance is represented by stars above each comparison (unpaired Student’s t-test; **** p < 0.0001 compared to the RNAi-FLUC control samples).

### Assessing protein-protein interactions (PPI) using a molecular docking consensus

The AaCHYMO midgut sequence (FASTA) was submitted to AlphaFold2 ^43^ via ColabFold ^44^ (v1.5.5; https://github.com/sokrypton/ColabFold) for modeling of its three-dimensional (3D) structure. The model, herein termed AaCHYMO, was predicted with options set as default, and structures were not relaxed using amber. Molecular docking was performed with three different software in a consensus approach for potential protein-protein interactions (PPI), assessing three proteins suggested as substrates: human serum albumin (PDB ID: 1E7H ^45^), human hemoglobin (PDB ID: 1GZX ^46^), and human neutralizing IgG (PDB ID: 1HZH ^47^). First, HDOCK ^48^, which employs a combination of template-based modeling and free docking, was used by submitting each pair of .pdb files (http://hdock.phys.hust.edu.cn/). Next, the pipeline for the extraction of predicted protein-protein interactions (PEPPI ^23^), which predicts the likelihood of sequence interactions, was employed by submitting each pair of FASTA sequences (https://zhanggroup.org/PEPPI/). Lastly, HADDOCK 2.4, which encodes information from predicted protein interfaces, was used with default parameters, while residue interaction restraints and patches were defined as random (https://rascar.science.uu.nl/haddock2.4/). The representative poses from HDOCK were selected based on docking scores and by visual inspection ^49^ of residues within feasible distances for potential PPI (∼5 Å) ^19,20^. The PyMOL software (v2.5.7) was used to analyze the results and generate images.

### Human Blood Protein Digestion Assays

AaCHYMO was incubated with human serum albumin (HSA) (Abcam #AB205808), hemoglobin (Hb) (MP Biomedicals, LLC. #55914), and immunoglobulin G (IgG) (Novus #NBP1-96779). In addition, for protein digestion comparison, human and bovine chymotrypsin were also incubated with the blood proteins. For these assays, a 10:1 (blood protein to protease) mass ratio was used. Reactions (50 μL) were carried out in 20 mM TRIS-HCl pH 7.2 and 10 mM CaCl_2_ at room temperature (25°C). To ensure that all activity assays were the same, prepared stocks that contained each blood protein, ultrapure water, and buffers (all added together, vortexed and centrifuged), then aliquoted into 1.6 mL microcentrifuge tubes at a volume of 46 μL (so that the final mass of the blood protein was 40 μg), followed by the addition of 4 μL of 1 μg/μL protease (4 μg of protease) to initiate digestion. All reactions were vortexed, centrifuged, and left at room temperature, withdrawing samples (6.1 μL to visualize 5 μg of total protein on a gel) at 10, 20, 40, and 60 min after the addition of the protease. A no-enzyme control (6.1 μL) was also withdrawn from the main stock of blood proteins without the proteases. All samples were treated with 3.4 μL of 6x SDS dye (sample buffer) and 13.9 μL ultrapure water and stored at −20°C until all samples were collected. Once collected, samples were thawed, vortexed, centrifuged, and incubated in a 95°C dry bath for 4 min. Samples were loaded onto 4-12% BIS-TRIS gels (Invitrogen #NP0322BOX) and ran at 175 volts for 40 min in the presence of 1x MES (Invitrogen #NP0002-02). The gel was then stained with Simply Blue Safe Stain (Invitrogen #LC6065) and destained with ultrapure water before photo documenting using the BioRad ChemiDoc MP Imaging System (Hercules, CA).

### Midgut Protein Extract Content Gel Analysis

With the stocks of FLUC and RNAi AaCHYMO midgut extracts (0.03 midgut/μL) used for the detection of chymotrypsin-like activity (see above), prepared 0.006 midgut equivalents for BIS-TRIS gel analysis. Samples were diluted with 20 mM TRIS-HCl pH 7.2 to a working stock concentration of 1.2×10^-3^ midgut/μL. From this diluted stock, 5 μL was removed and treated with 15 μl of ultrapure water and 3.4 μL of 6x SDS dye (sample buffer). Only the PBF samples were prepared for these experiments (3 h to 72 h PBF). Once all samples were prepared, they were vortexed, centrifuged, and incubated in a 95°C dry bath for 4 min. Samples were loaded onto 4-12% BIS-TRIS gels (Invitrogen #NP0321BOX) and ran at 175 volts for 40 min in the presence of 1x MES (Invitrogen #NP0002-02). The gel was then stained with Simply Blue Safe Stain (Invitrogen #LC6065) and destained with ultrapure water before photo documenting using the BioRad ChemiDoc MP Imaging System. The experiment was repeated in triplicate with different sets of mosquito tissue extracts. For quantification and statistical analysis, the most prominent protein band species were analyzed via ImageJ, and the pixel area of each determined. The pixel area was then plotted (in triplicate) and statistically analyzed by unpaired Student’s t-test with mean values ± SEM using GraphPad Prism version 10.4.0. Asterisks indicate significant differences (*p < 0.05; ns, not significant).

### Ecdysone nuclear receptor (EcR) knockdown, AaCHYMO protease expression, and activity determination

A gene encoding EcR was knocked down by the RNAi approaches described above, followed by processing of FLUC and EcR dsRNA injected mosquito tissues via sonication. Briefly, samples were removed from the −80°C freezer, thawed on ice, followed by quick vortex mixing (10 sec) and centrifugation (10 sec), then lysed using the dismembrator set at 25% amplitude (AMP) for 10 sec (3x total). To avoid overheating, samples were set on ice for about 30 sec, and the microtip was cleaned with ultrapure water and wiped dry with a Kimwipe to avoid cross-contamination. This was repeated for all samples. After lysing, all samples were then centrifuged at full speed in the VWR MicroStar 21 temperature-controlled centrifuge (14,800 rpm, 8°C) to pellet the midgut tissue away from the lysate. The lysates were then transferred to a new tube and immediately flash frozen in liquid nitrogen and stored at −80°C after use. To test activity, the MeO-Suc-Arg-Pro-Tyr-AMC commercially available substrate was used. A 10,000-fold dilution in activity assay buffer (20 mM TRIS-HCl pH 7.2 and 0.1% Tween 20) was prepared to test for activity (30 μL reactions). The final substrate concentration used was 25 μM to ensure linear regression activity during the run. A minimum of 3-6 replicates were performed for nearly all tissue extracts. Data was statistically analyzed by unpaired Student’s t-test with mean values ± SEM using GraphPad Prism. Asterisks indicate significant differences (**p < 0.05; ****p < 0.0001; ns, not significant).

### Fecundity Measurements of RNAi AaCHYMO Mosquitoes

The knockdown effect against AaCHYMO on fecundity was examined during the first gonotrophic cycle, as previously performed ^50^. Briefly, dsRNA-microinjected females were allowed to feed on bovine blood. Only fully engorged mosquitoes were chosen and allowed to lay eggs. Mosquitoes were individually transferred to a 15 mL vial containing wet oviposition paper at 48 hours after blood feeding and allowed to lay eggs for three days. Mosquitoes that were microinjected with dsRNA against FLUC were used as controls. Eggs were counted under a light microscope. Each dot represents the number of eggs deposited by individual RNAi-mosquitoes. Experiments were conducted from 3 biological cohorts using 20 mosquitoes (total of 60 mosquitoes). Unpaired Students *t*-test was used to determine whether a statistical difference in fecundity between RNAi-Fluc control group and RNAi-AaCHYMO is observed.

## ACKNOWELDGMENTS

Research reported in this publication was supported by the National Institute of General Medical Sciences (NIGMS) of the National Institutes of Health (NIH) under Award Number SC3GM116681 to AAR while at San José State University. The content is solely the responsibility of the authors and does not necessarily represent the official views of the National Institutes of Health.

## AUTHOR CONTRIBUTIONS

**Abigail G. Ramirez:** Conceptualization, Investigation, Formal Analysis

**Jun Isoe:** Conceptualization, Methodology, Validation, Formal Analysis, Investigation, Resources, Data Curation, Writing - Review & Editing, Visualization

**Mateus Sá M Serafim:** Conceptualization, Methodology, Validation, Formal Analysis, Investigation, Data Curation, Writing - Review & Editing, Visualization,

**Daniel Fong:** Investigation, Formal Analysis

**My Anh Le:** Investigation, Formal Analysis

**James T. Nguyen:** Formal Analysis, Writing - Review & Editing, Visualization

**Olive E. Burata:** Investigation

**Rachael M. Lucero:** Investigation

**Alberto A. Rascón, Jr.:** Conceptualization, Methodology, Validation, Formal Analysis, Investigation, Resources, Data Curation, Writing - Original Draft, Writing - Review & Editing, Visualization, Supervision, Project Administration, Funding Acquisition

## DATA AVAILABILITY

All data generated or analyzed during this study are included in this published article (and its Supplementary Information files) but may be made available from the corresponding author on reasonable request.

## SUPPLEMENTARY INFORMATION

**Supplementary Figure 1.**
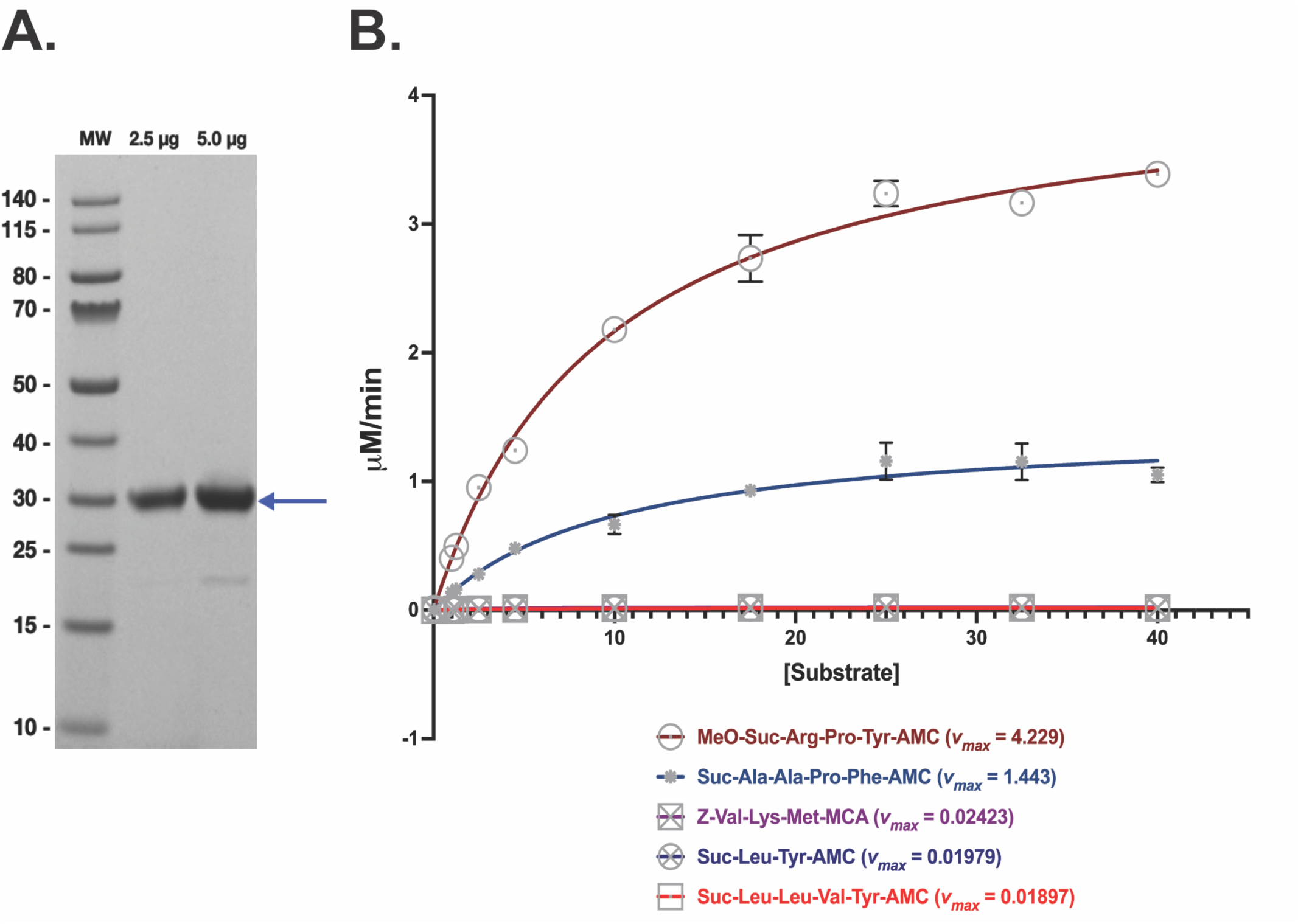
Activity assay determination of AaCHYMO under steady state conditions. **A.** Purified recombinant AaCHYMO (arrow). A total of 2.5 and 5 μg was loaded onto a NuPAGE^TM^ 4-12% BIS-TRIS gel (Invitrogen) to assess purity. MW represents the pre-stained PageRuler protein ladder in kilodaltons (kDa) (Thermo Scientific #26616). **B.** Michaelis-Menten plots of active recombinant AaCHYMO obtained from testing different commercially available chymotrypsin substrates (each is listed with their corresponding *v_max_*).

**Supplementary Figure 2.**
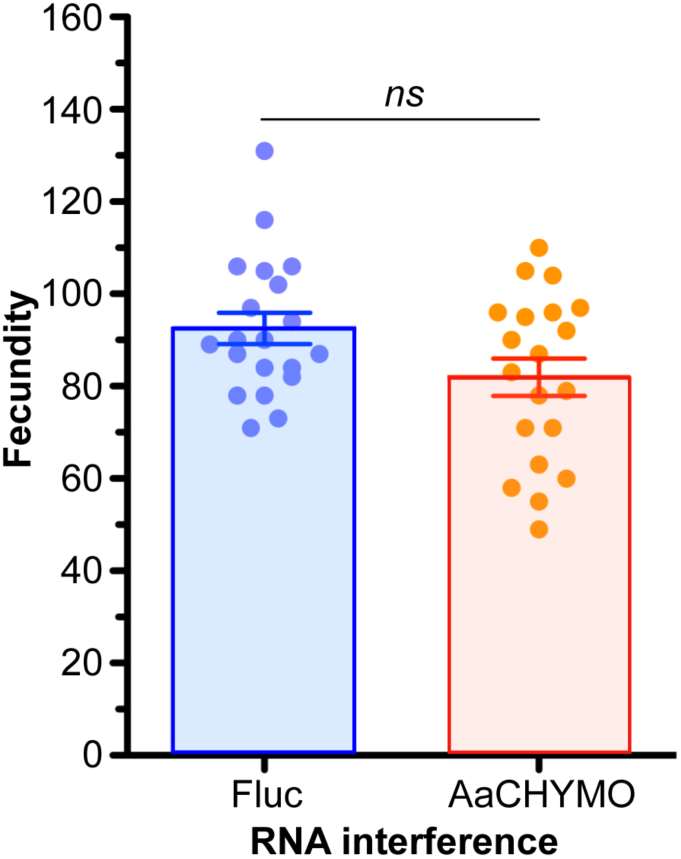
Effect of AaCHYMO dsRNA injection on fecundity in individual mosquitoes. No significant changes in the number of eggs oviposited are observed between FLUC and RNAi AaCHYMO mosquitoes (ns = no significance, p = 0.052). A total of 3 biological cohorts of 20 individual mosquitoes were chosen for fecundity analysis.

**Supplementary Table 1.** Commercially available and AaCHYMO-specific AMC substrates used.

| *Substrate (Commercially Available)* | *Company* | *Catalog #* | *Substrate (AaCHYMO-Specific Designed)* | *Company* | *Catalog #* |
| --- | --- | --- | --- | --- | --- |
| Suc-Ala-Ala-Pro-Phe-AMC | Biosynth | MAA-3114-V | Ac-Ala-His-Val-Phe-AMC | Biomatik | 1169025 |
| Suc-Ala-Ala-Phe-AMC | Biosynth | FS110519 | Ac-Ala-His-Leu-Phe-AMC | Biomatik | 1169029 |
| MeO-Suc-Arg-Pro-Tyr-AMC | Biosynth | SAP-3885-PI | Ac-Ala-His-Val-Tyr-AMC | Biomatik | 1169023 |
| Suc-Leu-Leu-Val-Tyr-AMC | Biosynth | MLL-3120-V | Ac-Ala-His-Leu-Tyr-AMC | Biomatik | 1169027 |
| Suc-Arg-Pro-Phe-His-Leu-Leu-Val-Tyr-MCA | Biosynth | MRP-3110-V | Ac-Ala-His-Thr-Trp-AMC | Biomatik | 1169037 |
| Suc-Leu-Tyr-AMC | Enzo Life Sciences | ALX-260-054-M005 | Ac-Ala-His-Asp-Trp-AMC | Biomatik | 1169039 |
| Suc-Ile-Ile-Trp-MCA | Biosynth | MIW-3150-V | Ac-Ala-His-Val-Trp-AMC | Biomatik | 1169038 |
| Ac-Ala-Asn-Trp-AMC | AdipoGen | AG-CP3-0037 | Ac-Ala-His-Val-Leu-AMC | Biomatik | 1169024 |
| Z-Leu-Leu-Leu-MCA | Biosynth | MLL-3177-V | Ac-Pro-His-Val-Met-AMC | Biomatik | 1169036 |
| Ac-Pro-Ala-Leu-AMC | AdipoGen | AG-CP3-0036 | Ac-Ala-His-Val-Met-AMC | Biomatik | 1169026 |
| Z-Gly-Gly-Leu-AMC | Thermo Scientific | J64796LB0 |  |  |  |
| Z-Val-Lys-Met-MCA | Biosynth | MVM-3156-V |  |  |  |
| Z-Arg-Arg-AMC | Biosynth | MRR-3123-V |  |  |  |
| Boc-Val-Leu-Lys-AMC | Biosynth | MVL-3104-V |  |  |  |

**Supplementary Table 2.**
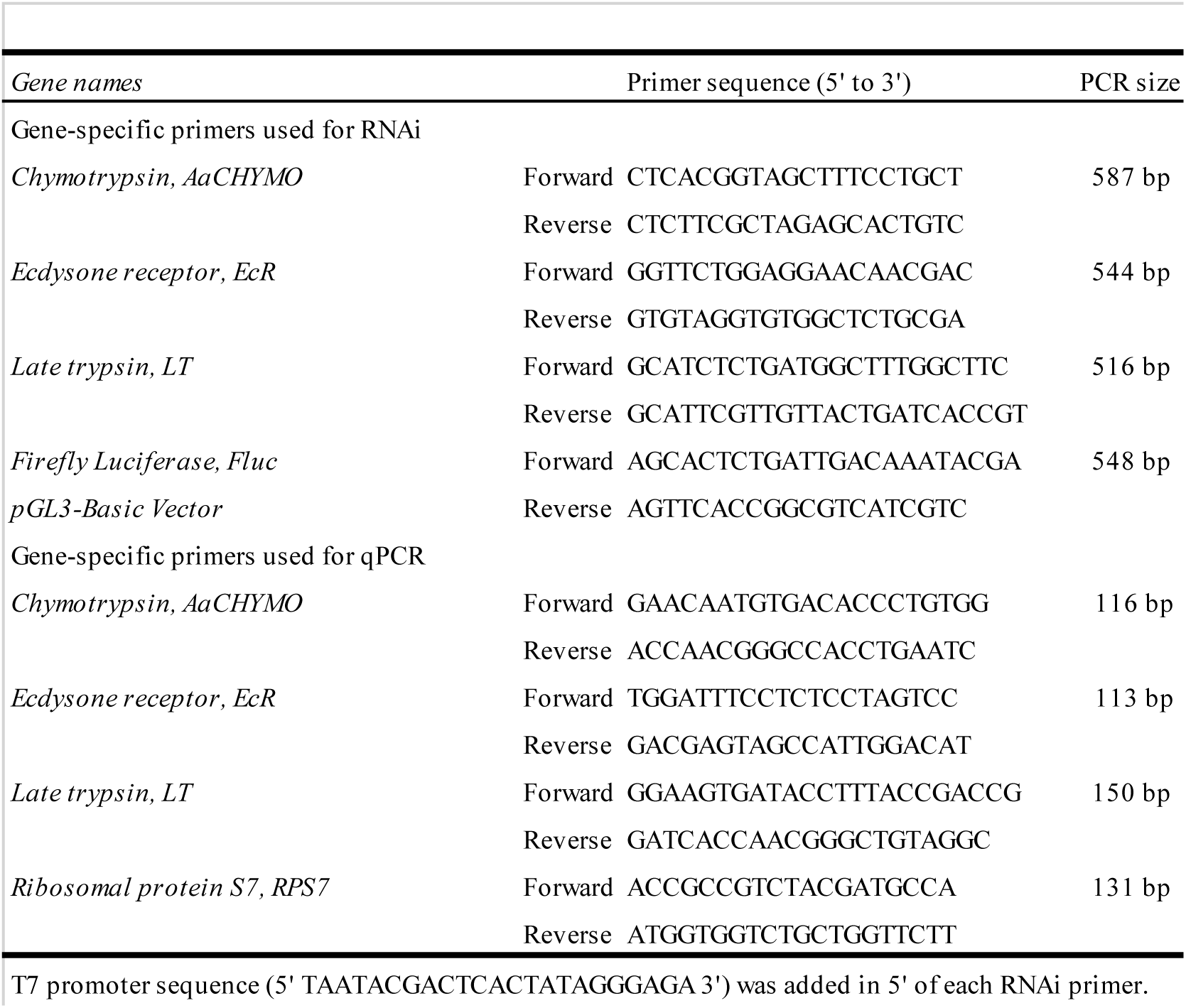
Gene-specific primers used for RNAi and qPCR.

